# Conditional Generation and Inpainting of Non-Coding RNA Sequences with Masked Discrete Diffusion

**DOI:** 10.64898/2026.09.17.752279

**Authors:** Utkarsh Upadhyay, Chuankai Dai, Julian Herold, Kengo Sato, Alexander Schug

## Abstract

Designing functional non-coding RNA (ncRNA) is fundamental to synthetic biology and RNA therapeutics, yet generative modelling for ncRNA has received far less attention than protein design. We present RNA-MDLM, a framework that extends Masked Discrete Language Models (MDLM) to the conditional generation and inpainting of ncRNA. We make two additions: first, conditioning on RNA-type representations from a pretrained RNA language model, and second, a modified classifier-free guidance scheme (Mod-CFG) that interpolates among conditional, unconditional, and random-sequence probabilities for better control. We also introduce REPAINT GAMES, a benchmark of seven structured masking tasks to probe a model’s performance on sequence patterns, structural motifs, and base-pairing per RNA type. Our model is trained on 4.6 million ncRNA sequences spanning six evaluable classes. It produces sequences whose composition and folding statistics closely match natural RNAs. Through extensive ablation studies, we find that the embedding-conditioned model achieves the best balance of structural fidelity, biological novelty, and inpainting accuracy, and its class label steers generation far more strongly than a plain label baseline. We further show that a model trained on a smaller, class-balanced subset can appear more realistic mainly by copying abundant natural sequences rather than learning their rules. We also benchmark against a masked-diffusion model and a family-specific VAE on ribozyme families, and find that a type-conditioned model like ours and a perfamily model are solving different tasks, which must be accounted for in a fair comparison. We will release the code and the trained models.

## 1 Introduction

Non-coding RNAs are one of the most versatile molecules in the cell, and their dysregulation can cause a wide range of human diseases. Transfer RNAs (tRNA) decode the genetic code, ribosomal RNAs (rRNA) form the catalytic core of the ribosome, small nuclear and nucleolar RNAs (snRNA, snoRNA) support splicing and rRNA modification, microRNAs (miRNA) silence genes post-transcriptionally (Bartel, 2018), and ribozymes catalyse reactions without proteins (Cech & Steitz, 2014). This diversity makes RNA an attractive therapeutic target to create mRNA vaccines, antisense oligonucleotides, engineered ribozymes (Damase et al., 2021; Sahin et al., 2014), and more. However, computational tools for designing new ncRNA sequences remain less developed than those for proteins.

Each RNA type has a distinct sequence-structure-function relationship: tRNAs adopt a cloverleaf formation, rRNAs form multi-stem conformations with long-range pairing, miRNAs fold into hairpin precursors, and ribozymes depend on precise active-site geometry. A model that generates plausible nucleotide strings is insufficient for design; it must produce reliable sequences that fold into the structures appropriate for their class. In protein engineering, generative models such as RFdiffusion (Watson et al., 2023), ProteinMPNN (Dauparas et al., 2022), and Chroma (Ingraham et al., 2023) have transformed design. RNA is catching up, with GenerRNA (Zhao et al., 2024), RNAdiffusion (Huang et al., 2024b), the masked-diffusion model EvoFlow-RNA (Patel et al., 2025), and large foundation models such as EVA (Huang et al., 2026).

**Figure 1:**
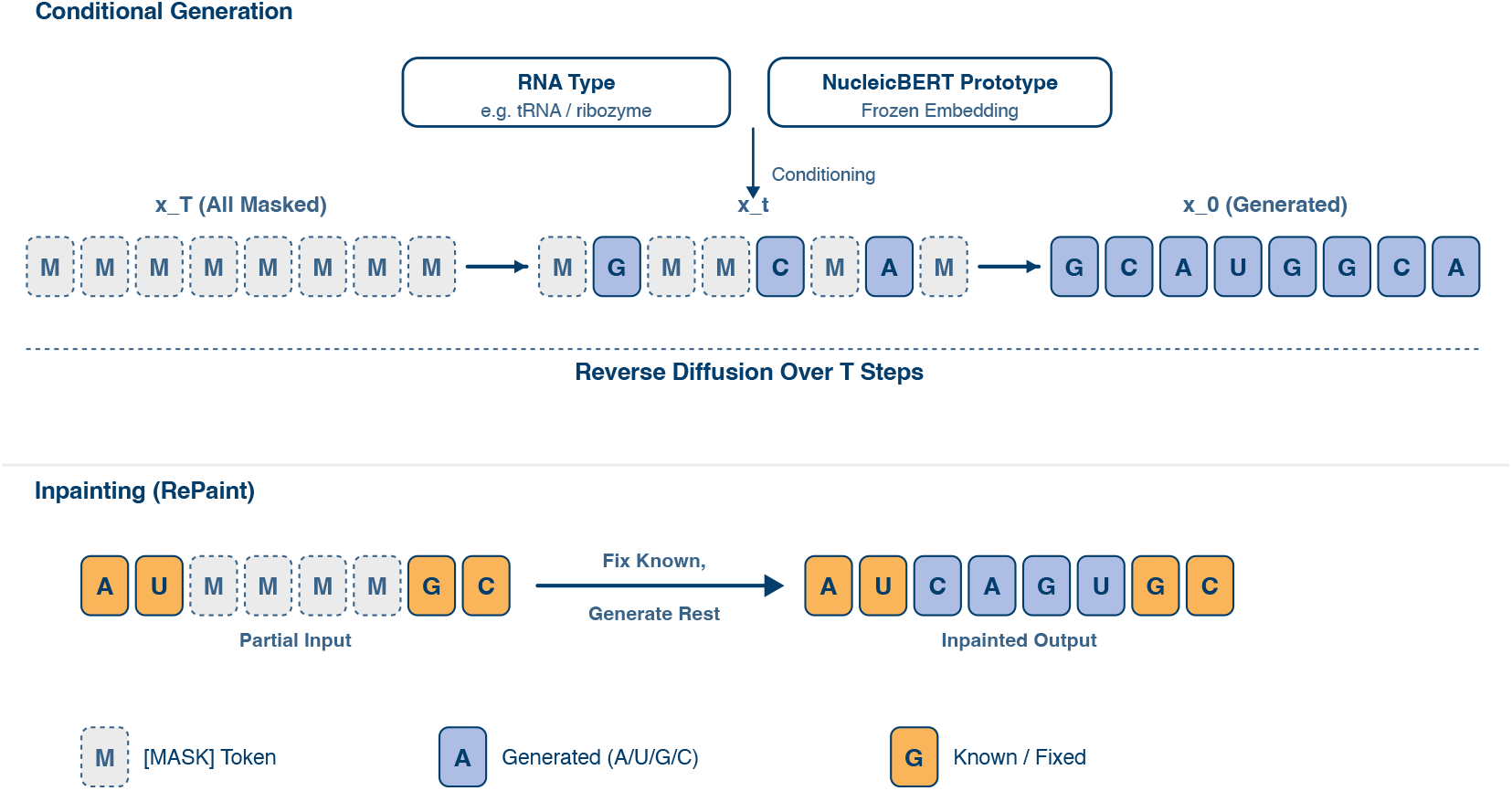
Overview of RNA-MDLM. Starting from a fully masked sequence *x_T_*, the DiT denoiser *p_θ_* iteratively unmasks tokens over *T* reverse-diffusion steps, conditioned on the RNA type. This conditioning can be provided through a simple label or through frozen NucleicBERT embeddings for each RNA type, to generate a novel sequence *x*_0_ that folds into a valid structure. The same model performs inpainting (bottom) where user-specified positions are held fixed (apricot) while masked positions are regenerated, enabling motif-constrained design.

For practical RNA design, producing a sequence that merely resembles a known RNA class is not sufficient. A useful generative model should preserve specified motifs or structural constraints while allowing the remaining sequence to change, and it should distinguish genuinely novel samples from sequences that closely reproduce examples seen during training. This creates a tension between fidelity and novelty: moving too far from the learned distribution may destroy class-specific constraints, whereas staying too close can make apparent generation quality difficult to distinguish from memorisation. A useful design model must therefore provide controllable generation while making clear whether biological fidelity reflects transferable sequence rules or proximity to examples already present in the data.

We find three major gaps in the practical design of *structured* ncRNA sequences. First, existing sequence-generative models provide limited systematic control over ncRNA type, often focusing instead on unconditional generation or individual families. Second, existing generative models do not naturally support *inpainting*, that is, keeping some known parts of the sequence fixed and generating the rest, a capability required in many RNA-engineering tasks where functional motifs or structural elements must be preserved while other regions are redesigned. Autoregressive models are particularly ill-suited to this setting because upstream nucleotides are sampled without access to downstream interaction partners, whereas downstream nucleotides use the already fixed upstream context. Third, there is no standard benchmark that systematically measures whether generative models capture class-specific sequence-to-structure relationships. This makes generative models difficult to compare under a common framework and forces individual studies to define their own evaluation protocols and metrics. Compounding all three is the size of sequence space: at length 100, more than 10^60^ sequences exist, so to navigate this space effectively, generation must be guided by strong biological priors.

Masked Diffusion Language Models (MDLM) (Sahoo et al., 2024) are absorbing-state discrete diffusion models. In these models, the forward process gradually replaces sequence tokens with a mask symbol, and the network is trained to reverse it by predicting the masked tokens over repeated denoising steps. Notably the bidirectional attention allows unresolved positions to be predicted from context on both sides and progressively refined within an increasingly complete sequence. This makes the same generative formulation suitable both for *de novo* generation from a fully masked sequence and for inpainting of partially specified RNA. Schiff et al. (2025) later derived classifier-free and classifier-based guidance for these models. We build on this model and adapt it to RNA. RNA-MDLM is trained from scratch as a Diffusion Transformer on 4.6M ncRNA sequences from RNAcentral (RNAcentral Consortium, 2021), annotated with RNA type, Rfam family, and Gene Ontology terms (we use only RNA type for conditioning here). The corpus has eight classes, but lncRNA (long non-coding RNA) and piRNA (piwi-interacting RNA) have too few sequences for evaluation, so we focus on the six prominent classes (tRNA, rRNA, miRNA, snRNA, snoRNA, ribozyme). *Our contributions:*

1. **A standardised dataset.** The field lacks a common dataset for multi-class ncRNA generation, so we curate 4.6 million type-annotated ncRNA sequences and train a single model that can generate any of the six evaluated RNA types.
2. **Embedding conditioning.** We replace the simple integer label conditioning with a frozen per-type representation from the pretrained NucleicBERT (Upadhyay et al., 2026) RNA language model. This conditioning gives the best overall balance of structural fidelity, low memorisation, and inpainting accuracy.
3. **Mod-CFG.** A modified classifier-free guidance scheme that interpolates among conditional, unconditional, and random-sequence predictions, adding two pseudo-classes to training so guidance has stable reference points even for the rarest types.
4. **Discrete RePaint and REPAINT GAMES.** We adapt RePaint (Lugmayr et al., 2022) to discrete absorbing-state diffusion for RNA inpainting, and introduce REPAINT GAMES, seven structured masking tasks that systematically probe arbitrary-context editing and class-specific sequence-structure constraints, providing a reproducible benchmark for comparing RNA generative models.

Beyond these components, we benchmark RNA-MDLM against a masked-diffusion model (EvoFlow-RNA) and a family-specific VAE (RfamGen) on two ribozyme families, and find that a type-conditioned generalist like RNA-MDLM and a per-family specialist are solving different tasks, which a fair comparison must account for. We further find that data curation changes what the model learns. A model trained on a smaller, class-balanced subset can appear more realistic mainly by copying abundant natural sequences, while clustering out near-duplicates has the opposite effect. A high fidelity score can therefore reflect either learning or copying, so we argue that generative RNA models should report novelty next to structural fidelity.

## 2 Related Work

### RNA generative models

Early RNA design used covariance models and stochastic context-free grammars (Nawrocki & Eddy, 2013), which describe families but do not generate novel sequences beyond a profile. Inverse-folding methods design a sequence for a target structure: RNAinverse (Hofacker et al., 1994) at the secondary-structure level, gRNAde (Joshi et al., 2025) and RiboDiffusion (Huang et al., 2024a) from 3D backbones. These are structure-constrained and cannot generate without a target. At the same time, there are generative models for RNA sequences: CodonBERT (Li et al., 2024) optimises mRNA (messenger RNA) codons; RfamGen (Sumi et al., 2024) designs sequences for a *single* family with a VAE that takes the family’s alignment, covariance model and consensus secondary structure as input; GenerRNA (Zhao et al., 2024) is an autoregressive model; RNAdiffusion (Huang et al., 2024b) does property-guided continuous latent diffusion; and EVA (Huang et al., 2026) is a large autoregressive foundation model. Closest to us, EvoFlow-RNA (Patel et al., 2025) also uses masked discrete diffusion, but builds on a pretrained BERT encoder and targets unconditional generation and motif scaffolding rather than explicit multiclass, type-conditional generation. RNA-MDLM is distinguished because of the following features: absorbing-state masked diffusion, per-type conditioning, controllable inpainting, and an inpainting benchmark, all trained from scratch. Appendix Figure 3 contrasts the two paradigms these models are built on, autoregressive generation and continuous-latent diffusion, and the limitations that motivate our masked discrete-diffusion approach.

### Discrete diffusion and guidance

D3PM (Austin et al., 2021) introduced discrete diffusion via structured transition matrices; SEDD (Lou et al., 2024) added score-entropy training; MDLM (Sahoo et al., 2024) unified these through a substitution parameterisation with an absorbing mask state and a tight variational bound. For biological sequence, DDSM (Avdeyev et al., 2023) applied Dirichlet diffusion to short DNA regulatory regions. Gradient-based steering does not transfer to discrete tokens; Schiff *et al*. (Schiff et al., 2025) derived discrete classifier-free and classifier-based guidance for absorbing-state models, building on classifier-free guidance (Ho & Salimans, 2021) and reparameterized discrete diffusion for text (Zheng et al., 2024). Our Mod-CFG adds a random-sequence reference term motivated by the class imbalance of RNA databases.

### Inpainting

RePaint (Lugmayr et al., 2022) inpaints with continuous DDPMs by overriding known pixels with a noised version of the ground truth and resampling to harmonise regions, without retraining. We adapt this to discrete absorbing-state diffusion, where known positions are noised through the mask token rather than additive Gaussian noise.

## 3 Methods

### 3.1 Model Architecture

RNA-MDLM uses a standard MDLM backbone. A Diffusion Transformer (DiT) (Peebles & Xie, 2023) with *N* =12 blocks, hidden dimension *d*=768, and conditioning dimension *d_c_*=768 (*∼*130M parameters), conditioned via adaptive layer normalisation. Full architecture, attention, position encoding, and single-nucleotide tokenisation (12 tokens) are given in Appendix A.3. The conditioning vector combines a timestep signal and a class signal, **c** = SiLU(**W***_t_ϕ*(*t*)) + **f**_class_(*y*), where *ϕ*(*t*) is a sinusoidal timestep embedding and *y* is the RNA type. We disabled the optional length term in all runs and control length at sampling time instead. We compare two choices of **f**_class_:

- **Label embedding** (baseline): a learnable table **E** *∈* R^(^*^C^*^+1)*×*^*^dc^* with a null class for CFG dropout.
- **NucleicBERT embedding.** NucleicBERT (Upadhyay et al., 2026) is a BERT-style model trained on *∼* 23 million RNA sequences. Even without using structural labels in pretraining, this model is able to separate RNA functional classes in embedding space. We extract a 1024-d representation per sequence (averaging hidden states over its 33 layers and over positions), average these within each RNA type, L2-normalise, and project through a learnable **W**_nb_ *∈* R^1024*×*^*^dc^*. We compute these representations once, offline, so training the diffusion model only updates the projection **W**_nb_ and never runs NucleicBERT itself. The null class maps to a zero vector, keeping CFG compatibility.

### 3.2 Modified Classifier-Free Guidance

Classifier-free guidance sharpens a conditional model by contrasting it with a class-agnostic reference and extrapolating away from that reference (Ho & Salimans, 2021; Schiff et al., 2025; Zheng et al., 2024). Standard CFG obtains this reference by randomly dropping the class label during training, giving the unconditional model *p_θ_*(*·|*∅∥), and combines the two predictions as log *p*_cfg_ *∝ γ* log *p_θ_*(*·|y*) + (1 *− γ*) log *p_θ_*(*·|*∅∥). This works when the reference is a neutral average over all classes. Under our class imbalance it is not: tRNA and rRNA are more than 95% of the sequences, so the dropout-trained *p_θ_*(*·|*∅∥) is dominated by them and, for a minority class such as snRNA, points toward tRNA/rRNA rather than toward a neutral average. Subtracting it then does little to isolate what makes the target class distinctive, and guidance is weak.

Mod-CFG replaces this with two explicit reference classes added to training. The first is a *pooled unconditional* class, built by sampling an equal number of sequences from every RNA type, so it is a balanced “average RNA” that removes the majority bias of the dropout reference. The second is a *random* nucleotide-string class, representing “no biological structure”. At each denoising step we form a per-token guidance score that boosts the conditional prediction and moves it away from *both* references,

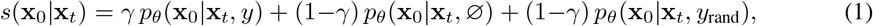

and sample the next token from its non-negative part, renormalised,

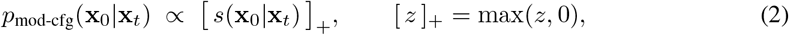

where a single weight *γ >* 1 sets the strength (we use *γ* = 5) and the two references share the complementary weight 1 *−γ*, exactly as the single reference does in standard CFG. Because 1 *−γ <* 0, the score *s* is not a probability distribution: it is an unnormalised vector whose entries may be negative, in the same way that standard CFG operates on an extrapolated (unnormalised) quantity. We never treat *s* as a probability. Sampling uses the same Gumbel-max categorical step as the rest of MDLM, which ranks tokens by their relative score and can never select a token with a non-positive weight; this realises the clamped, renormalised distribution of Eq. (2) exactly, without materialising the normalisation. Subtracting the balanced unconditional reference removes features common to RNA of any type, while subtracting the random reference removes generic nucleotide statistics, so what remains is specific to the requested class, and guidance stays informative even for the rarest types. A schematic is in Appendix Figure 4.

### 3.3 Discrete RePaint for RNA

Given **x**_0_ with known positions *K* and unknown positions *U*, inpainting proceeds as in Algorithm 1. At each reverse step, the model denoises the full sequence, then we override the known positions with samples from the forward process applied to the original tokens, which keeps them anchored while information flows into *U* through self-attention. Resampling (*R >* 1), jumping back *j* steps each round, harmonises the boundary between known and generated regions. Unless noted otherwise, we use *R* = 10, *j* = 10, *T* = 250. The only change from continuous RePaint is that we noise known positions by masking, 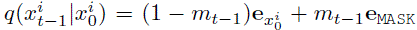, with no additive noise and no change to the trained model.

### 3.4 RePaint Games: A Systematic Inpainting Benchmark

REPAINT GAMES is a suite of seven structured masking tasks on real RNA, each a challenge with a clear success criterion rather than an arbitrary mask. **G1 Random Masking:** mask fractions *f ∈* {0.1, …, 0.9}, a general recovery measure. **G2 Contiguous Block:** mask one block at the 5’ end, centre, or 3’ end, isolating one– vs two-sided context. **G3 Motif Recovery:** using ViennaRNA structures, mask all loop or all stem positions. **G4 Flanking Context:** keep only terminal flanks (5– 30% each), reconstruct the interior. **G5 Interior Context:** keep a central segment (20–80%), extend outward. **G6 Cross-Type Conditioning:** 50% masking under correct, null, and wrong labels, testing label steering. **G7 Structure Preservation:** mask the 3’ (closing) side of every stem while keeping the 5’ side, probing Watson–Crick/wobble pairing learned from sequence alone. For each game, we report the mean recovery rate (fraction of masked positions correctly recovered) over 100 validation sequences, generating 4 samples each.

## 4 Experimental Setup

### Dataset

From RNAcentral (RNAcentral Consortium, 2021) we extract sequences of length 20– 512 nt that contain only standard nucleotides and that carry RNA type, Rfam family, and Gene Ontology terms as annotations. This filtering gives us 4,628,983 sequences, spanning eight RNA types of extremely different sizes like tRNA (2,445,367), rRNA (1,979,524), snoRNA (136,027), miRNA (41,766), ribozyme (18,425), snRNA (7,173), lncRNA (697), piRNA (4). We restrict evaluation to the six classes with *≥*1000 sequences. The imbalance (tRNA+rRNA *>*95%) is a property of the biology, not preprocessing. We keep 80% of the data for training and compute all metrics on the remaining data, which becomes the validation set. To probe redundancy we clustered the training data with CD-HIT-EST (Fu et al., 2012) at 90% and 80% identity (611,787 and 147,236 sequences). To disentangle data scale from imbalance, we built two class-balanced subsets: *500K* (50K per class + 150K random strings) and *50K* (10K each of five classes; it therefore has no snRNA). Details and length distributions are in Appendix A.6 and Figure 6.

### Training

All variants share the same DiT network (*d*=768, 12 blocks, 12 heads). We train them with AdamW (weight decay 0.1, gradient clipping 1.0) in bfloat16 and keep an EMA of the weights (decay 0.9999). The model with NucleicBERT embeddings uses a per-GPU batch of 64 on 32 nodes at learning rate 5 *×* 10*^−^*^4^. The other variants use a per-GPU batch of 300 on 16 GPUs at a learning rate 1.5 *×* 10*^−^*^4^. For each case, the checkpoint with the lowest validation loss was used for testing. The Subset 500K model is the one trained with Mod-CFG. It keeps every label (*p*_drop_ = 0) and includes the two pseudo-classes from Section 3.2. The other variants use standard guidance and drop the type label with probability *p*_drop_ = 0.1.

### Variants

We compare six models that share the same network and differ only in their training data or in how the RNA type is encoded. **Baseline** uses the full 4.6M dataset with a plain learned label (an integer index mapped to an embedding). **NucleicBERT** uses the same data but encodes the type with the frozen NucleicBERT prototype (Section 3.1). As a model name, “NucleicBERT” always means this diffusion model conditioned on NucleicBERT embeddings, not the NucleicBERT language model itself. **Subset 500K** and **Subset 50K** use the smaller class-balanced subsets, and **Cluster 80** and **Cluster 90** use the CD-HIT-clustered data. Subset 500K is also the one variant trained with Mod-CFG, since its data includes the random pseudo-class; all others use standard label guidance. Baseline and NucleicBERT differ only in the type encoding, so comparing them isolates the conditioning effect.

### Metrics

We report different metrics for the variety of tests. For composition, GC content and *k*-mer Jensen–Shannon divergence (*k ∈ {*2, 3, 4*}*) against the validation set. For structure, MFE per nucleotide, base-pair fraction, and ensemble diversity from ViennaRNA (Lorenz et al., 2011), against length-matched validation sequences. For novelty, the nearest-neighbour percent identity to the validation set, split into fractions above 95%, in the biological 60–90% band, and below 50% (natural Rfam variation sits near 73% (Kalvari et al., 2021; Upadhyay et al., 2026)). For inpainting, the REPAINT GAMES recovery rate.

## 5 Results

### 5.1 Sequence Composition and Novelty

Appendix Table 6 reports composition metrics for the NucleicBERT model against the held-out split. Generated GC content is close to the validation set for every type (miRNA 0.459 vs. 0.451; largest gaps are for ribozyme 4.5 and snoRNA 4.0 points), and *k*-mer JS divergences are small (2-mer 0.03–0.06), so local statistics are reproduced well. Crucially, the model achieves this realism while staying novel: the NucleicBERT model has mean identity 61–80% across types, with most sequences in the 60–90% band and no exact matches. Its only sizeable near-duplication is on rRNA, expected for such a conserved RNA type. By contrast, the Subset 500K model collapses onto a narrow region of rRNA space (94.7% mean identity), and the clustered models show essentially no near-duplication. Appendix Figure 7 shows the full per-variant comparison of *k*-mer divergence and identity categories.

### 5.2 Secondary Structure

Appendix Table 4 gives secondary-structure statistics for all six types and variants. The following findings are crucial. First, the NucleicBERT model matches natural structure most closely while staying novel: tRNA MFE/nt *−*0.337 (validation *−*0.369) and base-pair fraction 0.601 (validation 0.616); snoRNA MFE/nt *−*0.213 (validation *−*0.215). Second, Subset 500K attains the best raw energetics for tRNA/rRNA, but by near-duplication because 77.1% of its rRNA outputs are near-duplicates (Table 5), so the realism reflects reproduction of abundant rRNA rather than learned rules. Third, clustering keeps outputs novel but lowers structural realism (Cluster 90 tRNA MFE/nt *−*0.259 vs. *−*0.369); it means removing near-duplicates discards the examples that show which positions in a conserved motif may vary.

### 5.3 RePaint Games

Pooled over the seven games and the five classes present in every subset, NucleicBERT and Baseline lead at 0.53 overall, ahead of Subset 500K (0.49), Subset 50K (0.43), and the clustered models (0.36– 0.37); the per-game breakdown is in Appendix Table 7 and figures in Appendix A.11. Recovery falls smoothly with masking fraction (G1), and rRNA stays highest under heavy masking while miRNA falls fastest, likely because miRNA function depends on a short seed region at fixed positions that is hard to recover once masked. In motif recovery (G3), every variant recovers loops better than stems (*e.g.* 0.68 vs. 0.54 for NucleicBERT), because stems require long-range Watson–Crick pairing that local context cannot resolve. The most informative result is cross-type conditioning (G6), as we see that removing the correct label collapses NucleicBERT recovery from 0.69 to 0.25 (a steering gap of 0.44), whereas the label-embedding Baseline barely moves (0.68 to 0.59, gap 0.09). The Baseline’s higher recovery under the wrong label is not an advantage; it means the Baseline largely ignores the class label and reconstructs from sequence context alone, so its label carries little information, whereas NucleicBERT’s sharp drop shows its label genuinely drives generation. In structure preservation (G7), NucleicBERT fills the closing strand of stems at 0.93 (rRNA) and 0.86 (tRNA) recovery rates, from sequence alone with no structural supervision, surpassing the Baseline on every class.

#### Data scale vs. diversity

Data scale and diversity pull in different directions. Full-data models (0.53) and Subset 500K (0.49) are close on inpainting, while Subset 50K drops to 0.43. More data does not buy structural realism either: the full-data models are trained on roughly nine times as many sequences as Subset 500K, yet do not beat it on folding, and Subset 500K’s apparent edge there comes partly from near-duplication rather than better-learned rules. Clustering hurts inpainting most (0.36–0.37) even at comparable data size, because removing near-duplicates strips the small-perturbation examples that teach which positions tolerate variation, leaving a coarser class model. For editing applications, training on raw, unclustered data is therefore preferable, provided memorisation is monitored.

### 5.4 Comparison with Related Generative Models

Many contemporary RNA sequence generation models use different forms of conditioning and rarely provide any output-length control. This makes a plain unconditional comparison unfair. If we generate from each model without conditioning and compare folding statistics, the models produce different length distributions, and length alone drives most folding metrics. We therefore restrict the comparison to models whose output length we can set, and we generate sequences at the family’s natural median length. We use the two ribozyme families that RfamGen (Sumi et al., 2024) provides, glmS (*∼*150 nt) and Pistol (*∼*71 nt). We score each model on two axes: *fidelity*, the Infernal (Nawrocki & Eddy, 2013) covariance-model (CM) bit-score computed with `cmalign`, and *novelty*, the CM-independent identity to real family members, which neutralises RfamGen’s advantage of being scored by the CM it used as an input. We set the membership threshold at the 5th percentile of the real members’ bit-scores, so 95% of real members score above it, and we count a generated sequence as reaching the family if it scores at least this threshold. We generate 1000 sequences per model per family. RNA-MDLM is prompted with the *ribozyme* class label (not a family label) at the family’s length, EvoFlow-RNA (Patel et al., 2025) (33M) generates unconditionally at that length, and RfamGen uses a dedicated per-family model. Table 1 reports the outcome, and Appendix Figure 9 plots fidelity against novelty. For glmS, RNA-MDLM produces genuine family members that are *not* copies. It achieves a median bit-score of 65.0 against the members’ 81.5, with 75.8% reaching the membership band, at median identity 0.64 (members sit at 0.79). EvoFlow-RNA, which has no family conditioning, reaches neither family (0% in-family, negative median bit-scores). RfamGen sits at high fidelity and low novelty (bit-score 136.9, above the real members, identity 0.71), as expected for a per-family specialist scored by its own CM. On the fidelity–novelty plot the identity axis increases upward, so lower on that axis means more novel: genuine generation lies to the right of the member band (in-family) while staying low on identity (novel). RNA-MDLM is the only class-conditioned model to reach this region for glmS, clearing the membership band at the lowest identity to real members of any in-family model, so its sequences are members without being copies.

**Table 1:** Within-family conditional generation. Fidelity is the median Infernal CM bit-score (`cmalign`); novelty is the median identity to real members; in-family is the fraction reaching the members’ 5th-percentile bit-score band. 1000 generated sequences per model per family (RfamGen: 999 for Pistol; members: 943 glmS, 676 Pistol). RNA-MDLM is conditioned on the *ribozyme class*, not the family.

| Family | Model | Median bit-score | Median identity | In-family |
| --- | --- | --- | --- | --- |
| glmS (~150 nt) | Real members | 81.5 | 0.79 | 95.0% |
|  | RNA-MDLM | 65.0 | 0.64 | 75.8% |
|  | EvoFlow-RNA | -20.3 | 0.52 | 0.0% |
|  | RfamGen | 136.9 | 0.71 | 100% |
| Pistol (~71 nt) | Real members | 54.3 | 1.00 | 95.0% |
|  | RNA-MDLM | -14.6 | 0.53 | 0.6% |
|  | EvoFlow-RNA | -11.5 | 0.55 | 0.0% |
|  | RfamGen | 62.9 | 0.77 | 96.3% |

#### The Pistol result is steering, not coverage

On the Pistol family, RNA-MDLM appears to fail (0.6% in-family), but this is a category effect, not a coverage gap, and it is instructive. RNA-MDLM conditions on RNA *class*, not family. Within the ribozyme class Pistol is rare and length-crowded, it is 2.5% of the 18,425 ribozyme-class training sequences (glmS is 10.2%, 4*×* more), and at Pistol’s 60–80 nt length it makes only 5.1% of ribozymes while other families like hammerhead-II (39.9%) and Group II introns (37.9%) dominate (Appendix Figures 11–12). On comparing the 1000 generated sequences to the families, we find that 72.2% resemble hammerhead-II and only 4.8% resemble Pistol, matching its 5.1% training prevalence (Appendix Figure 13). At 71 nt, the class-conditioned model correctly samples the dominant hammerhead family and emits Pistol at its base rate, whereas glmS’s length (135–175 nt) happens to isolate it (76.8% of ribozymes there), so a class-plus-length prompt lands on glmS. Scoring a class-level generalist on family membership therefore measures the granularity of its conditioning, not the quality of its generation.

#### Caveats

Two points bound this comparison. First, both families are heavily represented in RNA-MDLM’s training (92% of Pistol and 45% of glmS members lie within 95% identity of a training sequence), so no clean held-out family benchmark exists here. The covariance model and the membership threshold both come from RfamGen’s own reference data, which favours RfamGen, so we also report the CM-independent novelty axis. Second, comparing a single class-conditioned generalist against per-family specialists is inherently a category mismatch. Rather than obscure this, we take it as the point that we should judge a type-conditioned model and a per-family model on what each was trained to do, and that’s why we opt for this within-family protocol.

#### Inpainting comparison

The comparison above is at the family level and favours the per-family specialist. Most tasks in the REPAINT GAMES suite don’t require a conditioning label; thus, we can compare with a model like EvoFlow-RNA. We use the same 100 validation sequences per game for every model and pool over the six RNA types and the six games EvoFlow-RNA can run; the cross-type game (G6) needs a class label, which EvoFlow-RNA lacks, so we omit it. Table 2 reports the outcome. RNA-MDLM leads overall (0.53 for the NucleicBERT variant and 0.52 for the label baseline, against 0.47), and the gap is largest on the structure-dependent games such as random masking (G1), motif recovery (G3), and closing-strand recovery (G7). On the two context-extension games, flanking (G4) and interior (G5), the overall numbers are similar, but the performance varies across RNA types. Per-type curves are in Appendix A.12.

**Table 2:** Inpainting comparison against EvoFlow-RNA on REPAINT GAMES, using the same masked sequences for every model. Values are mean recovery rate pooled over the six RNA types; Overall pools the six games shown. G6 (cross-type conditioning) is omitted because EvoFlow-RNA has no class label to steer with. The Baseline and NucleicBERT variants are statistically indistinguishable (paired Wilcoxon signed-rank *p* = 0.96; bootstrap 95% CI of the mean per-sequence recovery difference [*−*0.001, +0.004]; Cohen’s *d_z_* = 0.04), while both significantly outperform EvoFlow-RNA (*p <* 10*^−^*^13^, Cohen’s *d_z_* = 0.35).

| Model | G1 | G2 | G3 | G4 | G5 | G7 | Overall |
| --- | --- | --- | --- | --- | --- | --- | --- |
| Baseline | 0.61 | 0.54 | 0.60 | 0.38 | 0.44 | 0.70 | 0.52 |
| NucleicBERT | 0.62 | 0.55 | 0.61 | 0.39 | 0.45 | 0.69 | 0.53 |
| EvoFlow-RNA | 0.52 | 0.48 | 0.49 | 0.40 | 0.45 | 0.53 | 0.47 |

## 6 Discussion

### Realism can hide memorisation

We measure identity against held-out sequences, so a near-duplicate is a generated sequence almost identical to a natural one that was never used in training. The rate varies sharply across variants. The NucleicBERT model stays novel, with only about 10% near-duplicates even for rRNA. The training set for Subset 500K is built by taking 50K sequences for each of seven RNA types, repeating sequences for a type that has fewer than 50K, and adding 150K random strings. This balancing affects conserved and diverse classes very differently. rRNA is downsampled from about 2M sequences to 50K, but ribosomal RNA is so conserved that even those 50K are nearly identical. We observe that 77% of its rRNA outputs are near-duplicates, and the apparent realism reflects memorising abundant, redundant rRNA rather than learned rules. This observation is related to the conserved nature of an RNA type rather than its abundance. tRNA is equally abundant and also downsampled to 50K, but tRNA is diverse across isoacceptors and organisms. Hence, the same-sized sample still teaches general rules, and its Subset 500K outputs sit at 79.6% mean identity with no near-duplicates. Clustering is the opposite case, driving near-duplication to zero but at the cost of realism and inpainting. A model can thus score well on realism for two very different reasons, so any claim that a generative RNA model is data-efficient should come with an explicit novelty check.

### NucleicBERT conditioning as transfer learning

Using frozen embeddings as conditioning vectors is an indirect, inexpensive form of transfer learning (no gradients through NucleicBERT’s 32 layers). NucleicBERT was trained on *∼*23 million sequences without structural labels, yet its representations organise sequences into functional clusters, and the consistent gains, especially in structure preservation (G7) and cross-type steering (G6), show that this biological information transfers to generation. Since the conditioning vectors are recomputed prototypes, we can add new RNA classes or swap in improved pretrained models without retraining the backbone.

### Applications and limitations

The combination of conditional generation and inpainting supports RNA-engineering workflows. It can be used to generate specific RNA-type sequences and screen by folding energy, or replace a suboptimal region while preserving flanking context (*e.g.* transplanting a ribozyme core, or optimising a miRNA-precursor stem). Our current evaluation uses ViennaRNA thermodynamic predictions, which omit tertiary contacts and pseudoknots. Several directions extend this work, such as adding tertiary-structure prediction to the evaluation, validating designs through synthesis and functional assays, broadening REPAINT GAMES beyond 100 sequences per type, adding structure-aware training objectives, and better handling of class imbalance.

## 7 Conclusion

We presented RNA-MDLM, a masked-discrete-diffusion framework for conditional generation and inpainting of ncRNA. A single Diffusion Transformer generates six ncRNA classes with plausible secondary structures and easily recovers the masked parts of sequences across various masking tasks. The NucleicBERT-conditioned model gives the best balance of fidelity, novelty, and inpainting, and steers far more strongly than a label baseline; REPAINT GAMES offers a reusable test of the sequence-to-structure rules of RNA. Our ablations show that a model trained on a smaller, class-balanced subset can appear realistic mainly by copying abundant sequences, while clustering out near-duplicates has the opposite effect. A fidelity score alone can reflect either learning or copying, so generative RNA models should report novelty next to structural fidelity.

### AI USE STATEMENT

In this work, we used generative AI tools to assist with language editing of the manuscript and with routine coding support for analysis and plotting scripts. We did not use generative AI tools for research ideation, experimental design, or the generation of scientific results, which were produced by the authors. We have reviewed all AI-assisted work. The AI-assisted code was verified and tested by the authors, and all text was checked and edited by the authors. We take responsibility for the final content of this work, including all text, claims, and artifacts.

### Ethics statement

This work does not involve human subjects, personal data, or private information. All sequences are drawn from the public RNAcentral database, and all evaluation is performed in silico; no sequences were synthesised or functionally characterised. We note that generative models for functional RNA carry a potential dual-use dimension, since improved design of catalytic or regulatory RNA could in principle be misused. We consider the risk here limited because the model targets secondary-structure fidelity of common non-coding RNA classes rather than the design of pathogenic agents, and we report novelty and memorisation explicitly to avoid overstating its generative capability. We declare no conflicts of interest.

### Reproducibility statement

The model architecture is described in Section 3.1, the modified classifier-free guidance in Section 3.2, and the discrete RePaint procedure in Section 3.3 and Algorithm 1. Training configurations, dataset construction, clustering thresholds, and evaluation metrics are given in Section 4 and Appendix A.6, and the REPAINT GAMES tasks in Section 3.4. The within-family benchmarking protocol, including covariance-model scoring with Infernal and the novelty computation, is detailed in Section 5.4. All data derive from the public RNAcentral release, and complete per-type results appear in Appendix A. We will release the code, trained models, the REPAINT GAMES benchmark, and the benchmarking protocol as an anonymised repository with the submission.

## A Supplementary Results

### A.1 Model Overview

**Figure 2:**
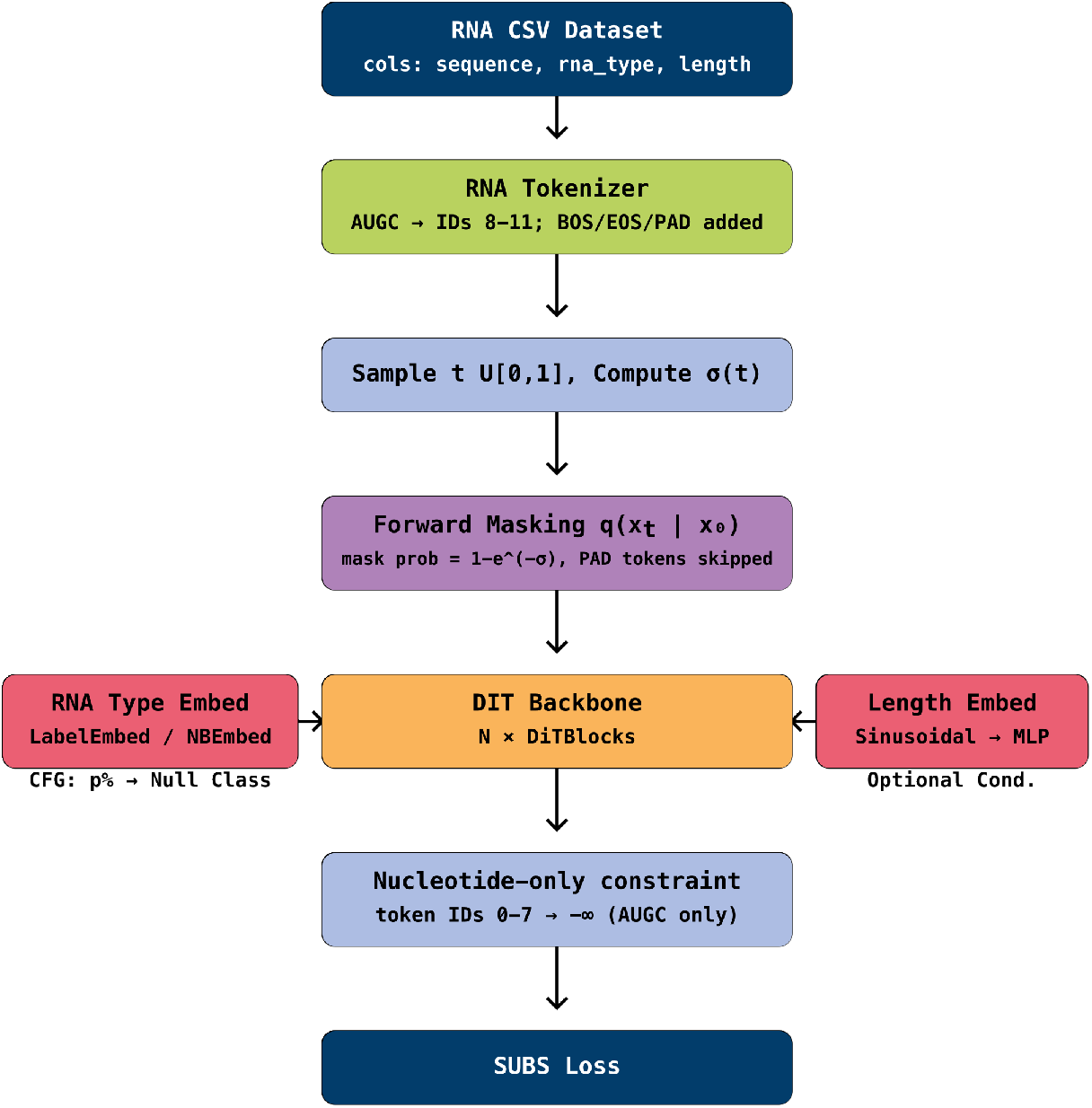
Overview of RNA-MDLM. RNA sequences are tokenised at single-nucleotide resolution and partially masked under the log-linear noise schedule. A Diffusion Transformer, conditioned on the RNA type through either a learned label or a frozen NucleicBERT prototype, is trained with the SUBS objective to predict the masked tokens. At sampling time the same model generates a full sequence of a chosen type from a fully masked input.

### A.2 Background: Masked Discrete Diffusion

Our model is built on MDLM (Sahoo et al., 2024); below is a brief explanation of how these models work.

Let **x**_0_ *∈* {1, …, *K*}*^L^* be a sequence of length *L* over a vocabulary of size *K*. The forward process independently replaces each token with a `[MASK]` token at a rate set by a continuous-time schedule *σ*(*t*), *t ∈* [0, 1]:

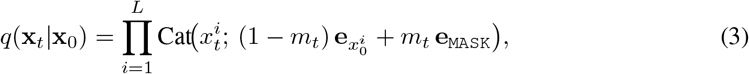

where *m_t_* = 1 *−* exp(*−σ*(*t*)) is the probability that a token has been masked by time *t*, and **e***_k_* is the one-hot vector for token *k*. At *t*=0 the sequence is clean, and as *t* grows more tokens become `[MASK]`. The reverse (denoising) model *p_θ_*(**x**_0_*|***x***_t_, t*) reads the partially masked sequence and predicts the original nucleotide at each position. The substitution (SUBS) parameterisation (Sahoo et al., 2024) adds two constraints. *Carry-over unmasking*: a revealed position is copied forward unchanged, so the model only predicts positions that are still masked. *Zero masking probability*: the model cannot output `[MASK]`, so a revealed position stays revealed. Together these make denoising *monotonic* which means over the reverse steps the masked set only shrinks, and a token is never remasked once committed. Under these constraints the objective reduces to masked-token prediction losses evaluated only at masked positions:

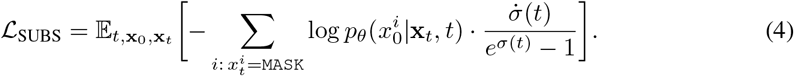

The factor 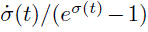 weights this loss toward intermediate noise levels, where the sequence is partly masked and the model must use real context to fill the gaps; this is where long-range structural patterns are learned. For our case we use the log-linear schedule, *σ*(*t*) = *−* log(1 *−* (1 *− ɛ*)*t*) with *ɛ* = 10*^−^*^3^, under which the masking probability from Eq. (3) becomes *m_t_* = (1 *− ɛ*)*t*, linear in *t*. To generate, we start from a fully masked sequence and run the reverse model for *T* steps until the sequence is complete. Appendix Figure 2 gives an overview, and the guided sampling loop is in Appendix Figure 4.

### A.3 Model Architecture Details

We use a Diffusion Transformer (DiT) (Peebles & Xie, 2023) as the denoising network, with adaptive layer normalisation (AdaLN-Zero) so the conditioning vector rescales and shifts the normalisation statistics at every layer. Each block uses multi-head self-attention (Vaswani et al., 2017) with Flash Attention (Dao et al., 2022) and rotary position embeddings (Su et al., 2024), which encode relative positions so the model recognises motifs such as stem-loops regardless of absolute location. The default configuration has *N* =12 transformer blocks, a hidden dimension *d*=768, 12 attention heads, and a conditioning dimension *d_c_*=768, for about 130M parameters. The RNA sequences are tokenised at a single-nucleotide resolution, which gives us the following 12 tokens: four nucleotides A, U, G, C; a `[MASK]` token for the absorbing state; and seven special tokens (`[CLS]/[SEP]/[BOS]/[EOS]/[PAD]/[RESERVED]/[UNK]`).

### A.4 Generation Paradigms

**Figure 3:**
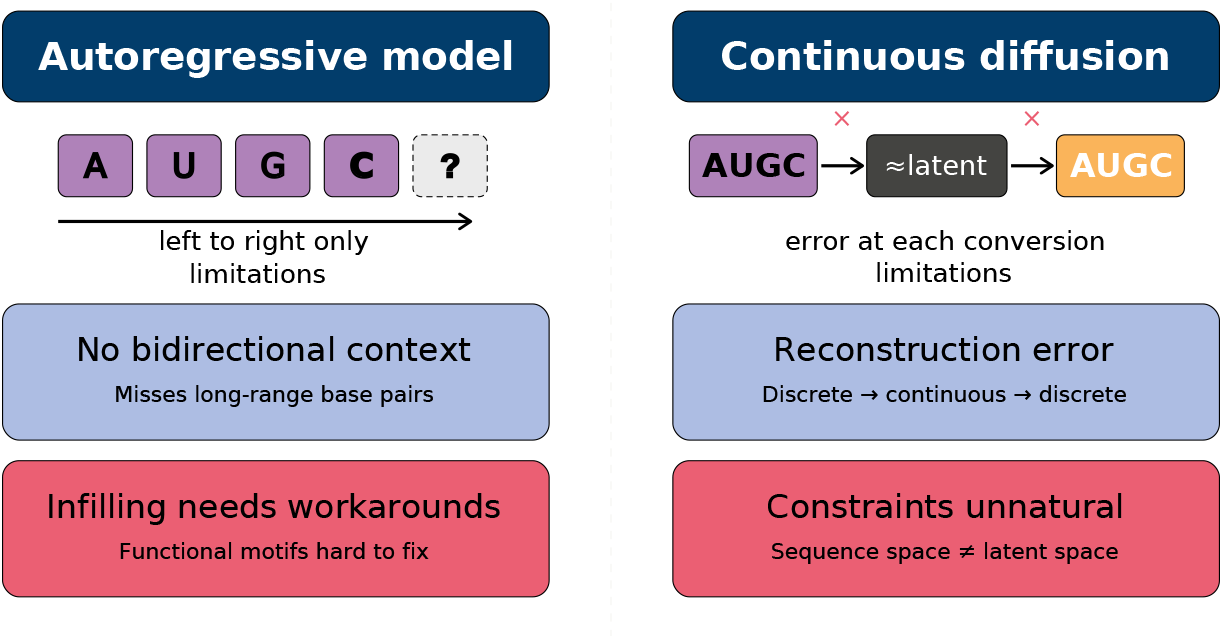
Two common paradigms for RNA generation and their limitations. Autoregressive models generate left to right, which makes bidirectional infilling less natural and can make long-range base pairs harder to exploit. Continuous diffusion moves between sequence and latent space, which can introduce reconstruction error and makes hard sequence constraints awkward to impose. RNA-MDLM uses masked discrete diffusion, which sidesteps both.

### A.5 Sampling Schematics

**Figure 4:**
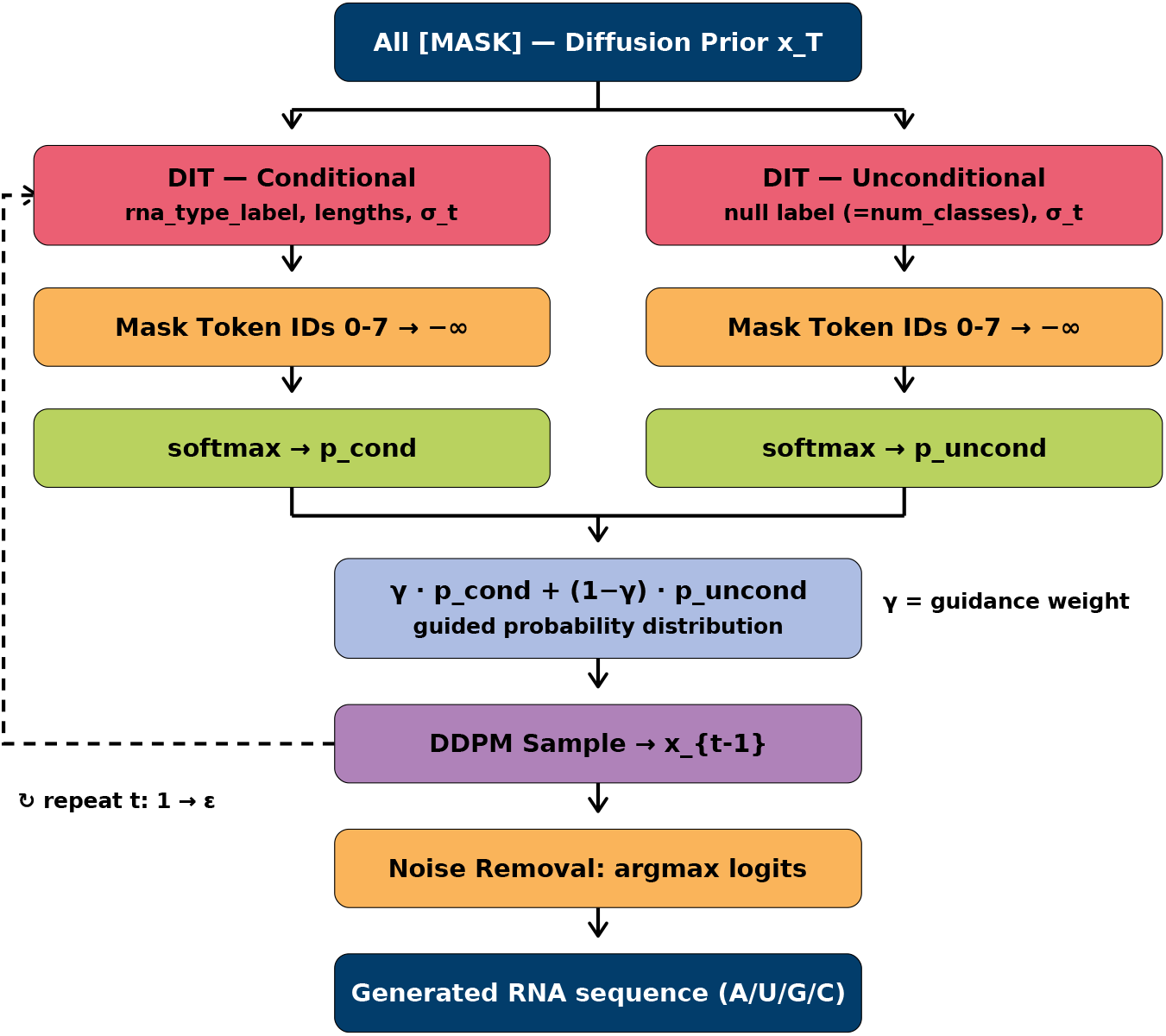
Classifier-free guided sampling in RNA-MDLM. At each step the conditional prediction is combined with the class-agnostic reference(s) and used to sample the next partially unmasked sequence.

**Figure 5:**
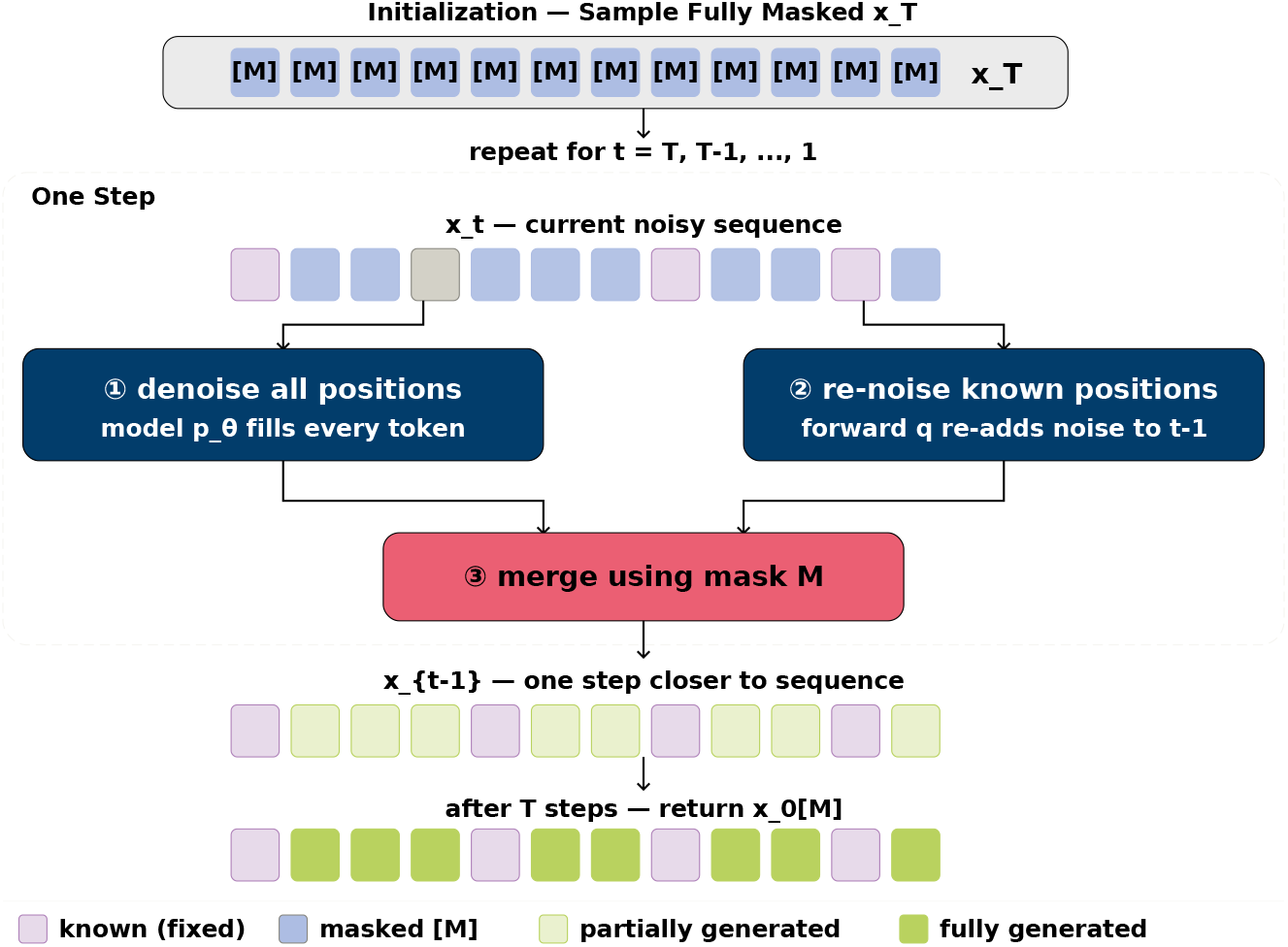
Discrete RePaint for RNA inpainting. Known positions are re-masked by the forward process and overridden at each reverse step; resampling harmonises the generated region with the fixed context.

**Algorithm 1.**
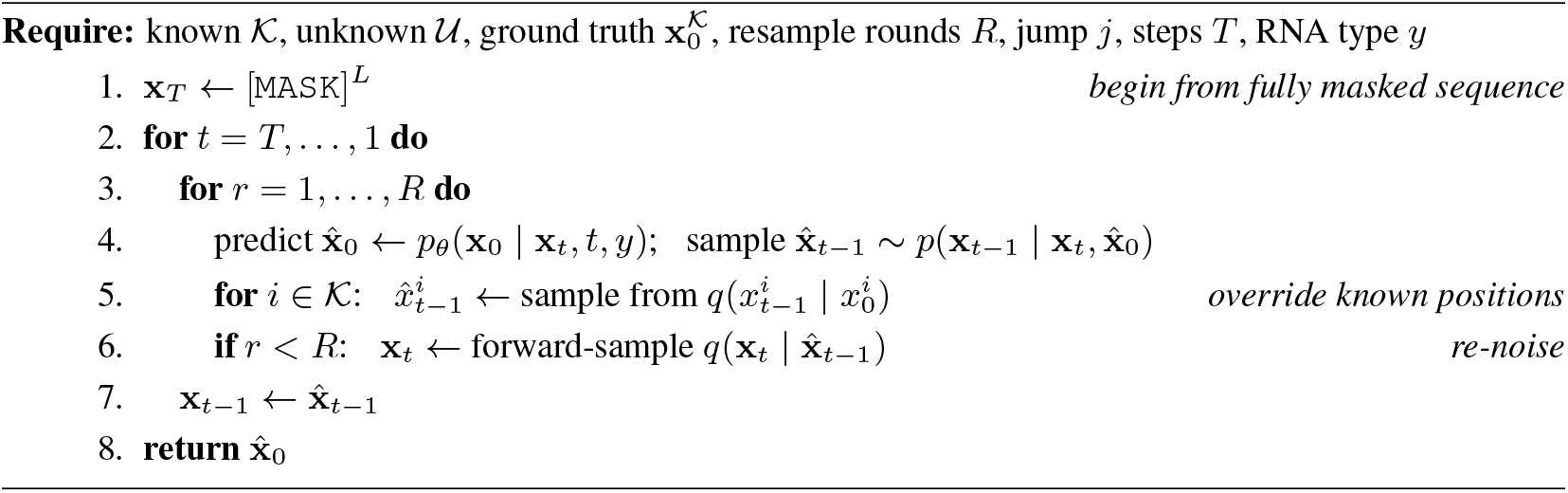
Discrete RePaint for RNA Inpainting.

### A.6 Dataset Construction

**Figure 6:**
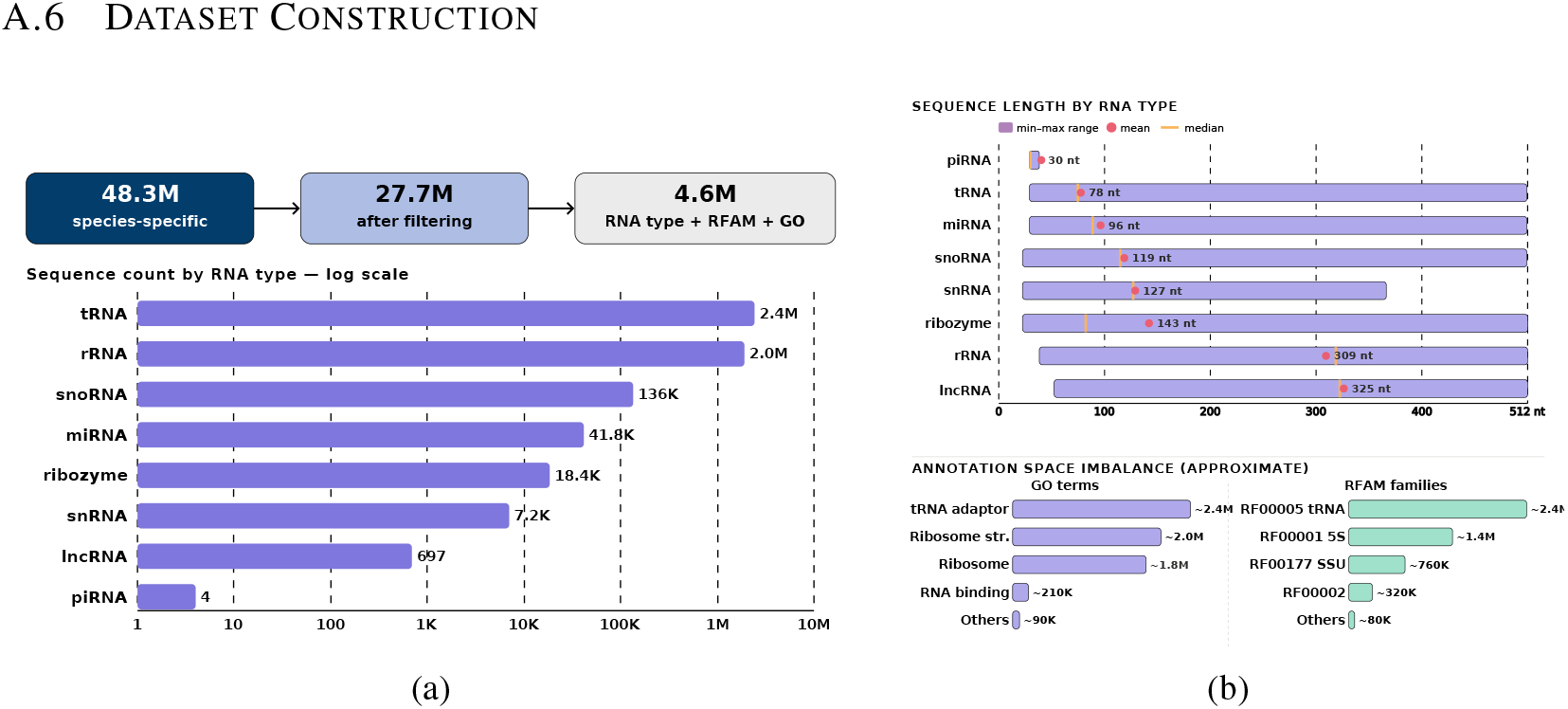
Dataset construction. (a) Filtering funnel from the raw RNAcentral download to the 4.6M annotated corpus (per-class counts, log scale). (b) Sequence-length distribution.

**Table 3:**
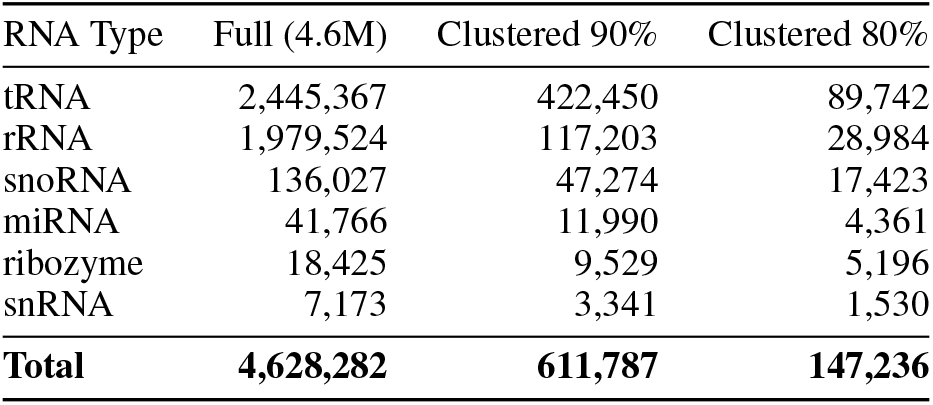
RNA type distribution across training datasets for the six evaluated classes. The Full column excludes lncRNA (697) and piRNA (4).

| RNA Type | Full (4.6M) | Clustered 90% | Clustered 80% |
| --- | --- | --- | --- |
| tRNA | 2,445,367 | 422,450 | 89,742 |
| rRNA | 1,979,524 | 117,203 | 28,984 |
| snoRNA | 136,027 | 47,274 | 17,423 |
| miRNA | 41,766 | 11,990 | 4,361 |
| ribozyme | 18,425 | 9,529 | 5,196 |
| snRNA | 7,173 | 3,341 | 1,530 |
| **Total** | **4,628,282** | **611,787** | **147,236** |

### A.7 Complete Secondary Structure Results

Table 4 reports metrics for all six RNA types and every variant as mean *±* standard deviation (over 1000 generated sequences per type, and over the length-matched validation set for the reference row).

**Table 4:** Secondary-structure metrics (mean *±* s.d.) for all six RNA types and variants. A dash denotes a configuration not evaluated for that type (the 50K subset has no snRNA).

| Model | MFE/nt | BP | ED |
| --- | --- | --- | --- |
| **tRNA (75 nt)** |  |  |  |
| Validation | $-0.369 \pm 0.08$ | $0.616 \pm 0.06$ | $11.35 \pm 6.1$ |
| Baseline | $-0.309 \pm 0.07$ | $0.583 \pm 0.07$ | $13.55 \pm 6.8$ |
| NucleicBERT | $-0.337 \pm 0.07$ | $0.601 \pm 0.06$ | $12.14 \pm 6.5$ |
| Subset 500K | $-0.391 \pm 0.06$ | $0.623 \pm 0.06$ | $9.94 \pm 5.9$ |
| Subset 50K | $-0.307 \pm 0.08$ | $0.592 \pm 0.07$ | $14.93 \pm 7.0$ |
| Cluster 80 | $-0.264 \pm 0.08$ | $0.563 \pm 0.08$ | $13.95 \pm 6.6$ |
| Cluster 90 | $-0.259 \pm 0.07$ | $0.564 \pm 0.08$ | $14.35 \pm 7.0$ |
| **rRNA (300 nt)** |  |  |  |
| Validation | $-0.413 \pm 0.05$ | $0.640 \pm 0.03$ | $63.97 \pm 20.6$ |
| Baseline | $-0.351 \pm 0.06$ | $0.640 \pm 0.04$ | $46.47 \pm 18.5$ |
| NucleicBERT | $-0.356 \pm 0.06$ | $0.641 \pm 0.04$ | $46.07 \pm 18.2$ |

Table 4 (continued)
| Model | MFE/nt | BP | ED |
| --- | --- | --- | --- |
| Subset 500K | $-0.418 \pm 0.04$ | $0.644 \pm 0.03$ | $62.27 \pm 18.3$ |
| Subset 50K | $-0.350 \pm 0.06$ | $0.637 \pm 0.04$ | $54.96 \pm 19.1$ |
| Cluster 80 | $-0.313 \pm 0.05$ | $0.601 \pm 0.04$ | $65.41 \pm 18.1$ |
| Cluster 90 | $-0.305 \pm 0.06$ | $0.608 \pm 0.05$ | $63.08 \pm 17.8$ |
| **miRNA (80 nt)** |  |  |  |
| Validation | $-0.439 \pm 0.10$ | $0.722 \pm 0.07$ | $5.90 \pm 3.6$ |
| Baseline | $-0.353 \pm 0.11$ | $0.644 \pm 0.09$ | $9.55 \pm 6.9$ |
| NucleicBERT | $-0.379 \pm 0.10$ | $0.675 \pm 0.09$ | $7.93 \pm 5.6$ |
| Subset 500K | $-0.395 \pm 0.08$ | $0.707 \pm 0.07$ | $7.17 \pm 4.2$ |
| Subset 50K | $-0.235 \pm 0.08$ | $0.579 \pm 0.09$ | $12.80 \pm 6.9$ |
| Cluster 80 | $-0.246 \pm 0.07$ | $0.575 \pm 0.08$ | $15.23 \pm 7.3$ |
| Cluster 90 | $-0.231 \pm 0.07$ | $0.565 \pm 0.08$ | $16.78 \pm 7.6$ |
| **ribozyme (80 nt)** |  |  |  |
| Validation | $-0.406 \pm 0.12$ | $0.644 \pm 0.08$ | $8.26 \pm 5.2$ |
| Baseline | $-0.342 \pm 0.11$ | $0.608 \pm 0.08$ | $13.04 \pm 7.3$ |
| NucleicBERT | $-0.327 \pm 0.10$ | $0.607 \pm 0.08$ | $12.47 \pm 6.5$ |
| Subset 500K | $-0.362 \pm 0.11$ | $0.678 \pm 0.05$ | $6.89 \pm 4.1$ |
| Subset 50K | $-0.306 \pm 0.10$ | $0.586 \pm 0.07$ | $16.22 \pm 7.7$ |
| Cluster 80 | $-0.350 \pm 0.10$ | $0.609 \pm 0.07$ | $13.41 \pm 6.8$ |
| Cluster 90 | $-0.269 \pm 0.07$ | $0.529 \pm 0.08$ | $15.63 \pm 7.2$ |
| **snRNA (120 nt)** |  |  |  |
| Validation | $-0.246 \pm 0.05$ | $0.558 \pm 0.06$ | $22.77 \pm 9.5$ |
| Baseline | $-0.233 \pm 0.06$ | $0.539 \pm 0.07$ | $24.47 \pm 9.1$ |
| NucleicBERT | $-0.221 \pm 0.05$ | $0.541 \pm 0.06$ | $24.08 \pm 9.6$ |
| Subset 500K | $-0.267 \pm 0.06$ | $0.568 \pm 0.06$ | $22.28 \pm 9.9$ |
| Subset 50K | – | – | – |
| Cluster 80 | $-0.228 \pm 0.04$ | $0.544 \pm 0.06$ | $25.85 \pm 9.2$ |
| Cluster 90 | $-0.172 \pm 0.05$ | $0.530 \pm 0.07$ | $26.09 \pm 9.3$ |
| **snoRNA (115 nt)** |  |  |  |
| Validation | $-0.215 \pm 0.07$ | $0.544 \pm 0.08$ | $21.94 \pm 8.9$ |
| Baseline | $-0.226 \pm 0.08$ | $0.550 \pm 0.08$ | $23.88 \pm 9.2$ |
| NucleicBERT | $-0.213 \pm 0.07$ | $0.549 \pm 0.07$ | $23.52 \pm 9.4$ |
| Subset 500K | $-0.279 \pm 0.07$ | $0.589 \pm 0.07$ | $22.17 \pm 9.8$ |
| Subset 50K | $-0.221 \pm 0.07$ | $0.553 \pm 0.07$ | $23.66 \pm 9.8$ |
| Cluster 80 | $-0.233 \pm 0.08$ | $0.543 \pm 0.08$ | $23.41 \pm 9.4$ |
| Cluster 90 | $-0.220 \pm 0.07$ | $0.564 \pm 0.07$ | $24.89 \pm 10.0$ |

### A.8 Novelty and Memorisation by Variant

**Table 5:**
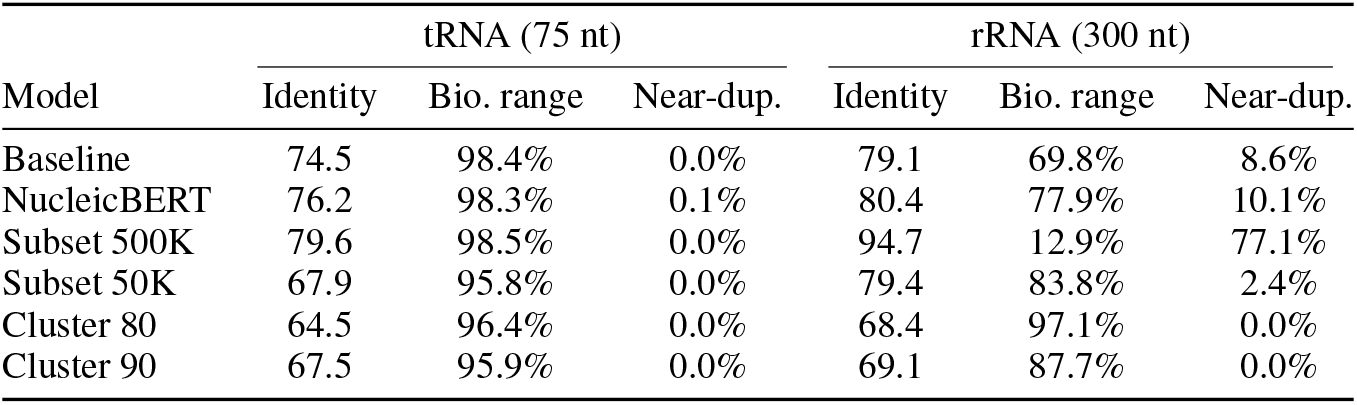
Novelty by variant for the two most abundant RNA types, against held-out validation. Mean identity is to the nearest validation neighbour; biological range is 60–90% (typical Rfam identity *≈* 73%); Near-dup. is the fraction above 95%.

| Model | tRNA (75 nt) |  |  | rRNA (300 nt) |  |  |
| --- | --- | --- | --- | --- | --- | --- |
|  | Identity | Bio. range | Near-dup. | Identity | Bio. range | Near-dup. |
| Baseline | 74.5 | 98.4% | 0.0% | 79.1 | 69.8% | 8.6% |
| NucleicBERT | 76.2 | 98.3% | 0.1% | 80.4 | 77.9% | 10.1% |
| Subset 500K | 79.6 | 98.5% | 0.0% | 94.7 | 12.9% | 77.1% |
| Subset 50K | 67.9 | 95.8% | 0.0% | 79.4 | 83.8% | 2.4% |
| Cluster 80 | 64.5 | 96.4% | 0.0% | 68.4 | 97.1% | 0.0% |
| Cluster 90 | 67.5 | 95.9% | 0.0% | 69.1 | 87.7% | 0.0% |

### A.9 Sequence Composition and Structure Deviation Figures

**Figure 7:**
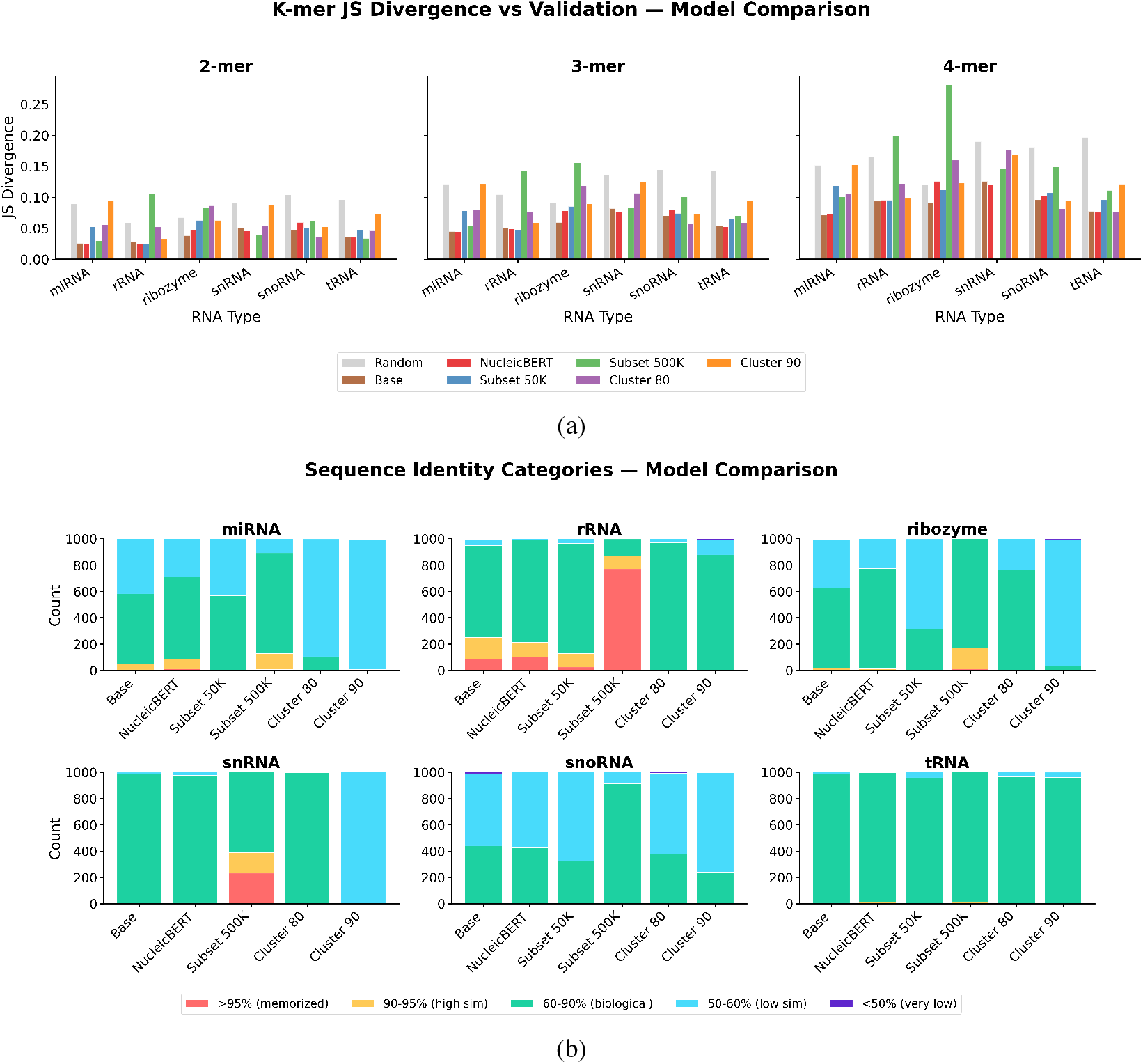
Sequence composition and novelty across the six variants. (a) *k*-mer JS divergence (*k ∈ {*2, 3, 4*}*) per type, with a random-sequence baseline for scale (lower is better). (b) Nearest-neighbour identity categories.

**Figure 8:**
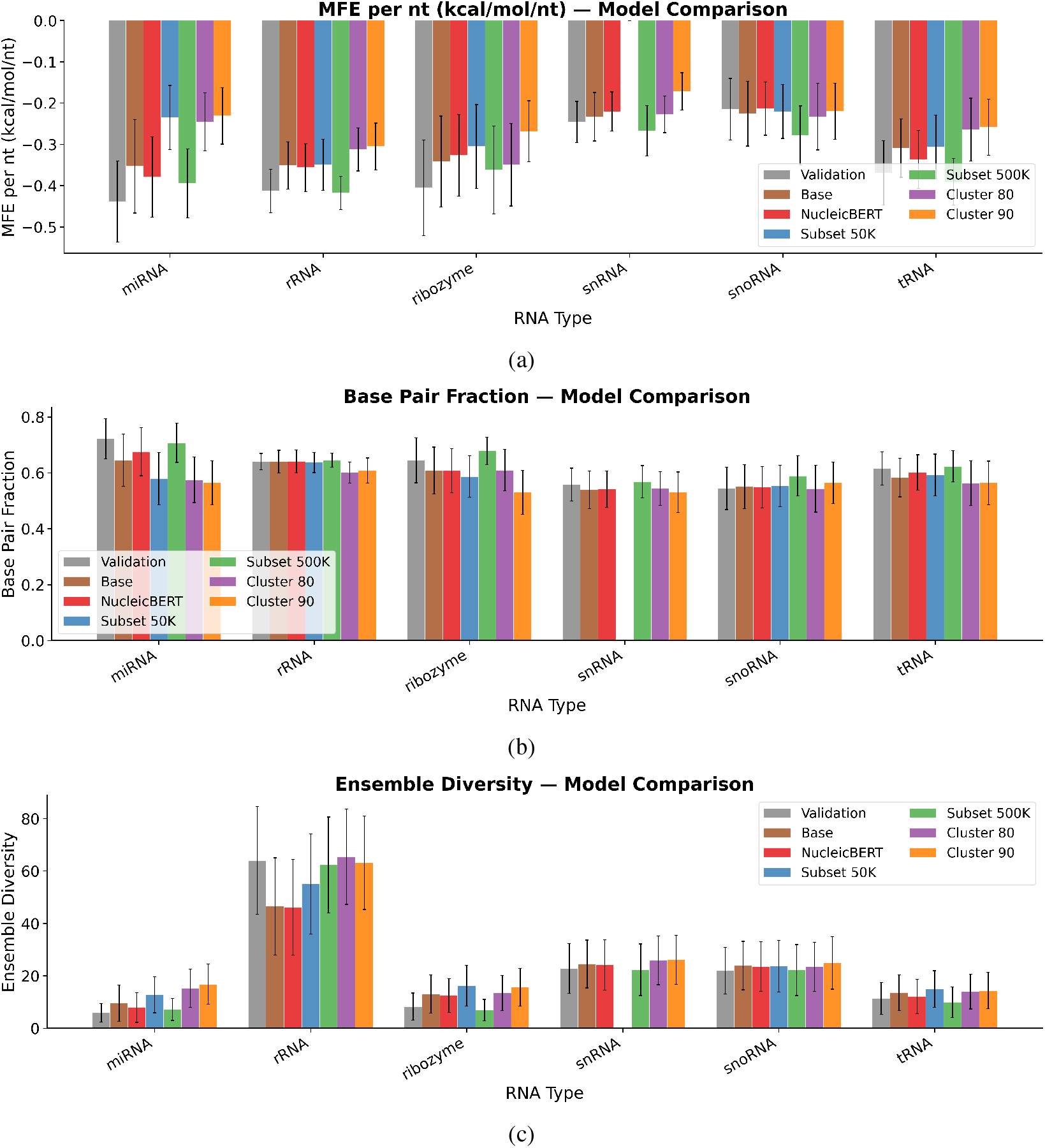
Secondary-structure statistics of generated sequences vs. the length-matched validation set, by RNA type and variant. (a) MFE per nucleotide. (b) Base-pair fraction. (c) Ensemble diversity.

**Table 6:** Compositional analysis of generated sequences (1000 samples per type, NucleicBERT model), against the held-out validation split.

| RNA Type | GC (Gen) | GC (Val) | 2-mer JS | 3-mer JS | 4-mer JS | Identity (%) |
| --- | --- | --- | --- | --- | --- | --- |
| tRNA | 0.543 | 0.567 | 0.035 | 0.052 | 0.075 | 76.2 |
| rRNA | 0.513 | 0.527 | 0.025 | 0.049 | 0.095 | 80.4 |
| miRNA | 0.459 | 0.451 | 0.025 | 0.044 | 0.073 | 70.7 |
| ribozyme | 0.501 | 0.546 | 0.047 | 0.078 | 0.126 | 66.3 |
| snRNA | 0.432 | 0.445 | 0.046 | 0.076 | 0.120 | 75.1 |
| snoRNA | 0.406 | 0.446 | 0.060 | 0.080 | 0.101 | 61.0 |

### A.10 Benchmarking Diagnostics

**Figure 9:**
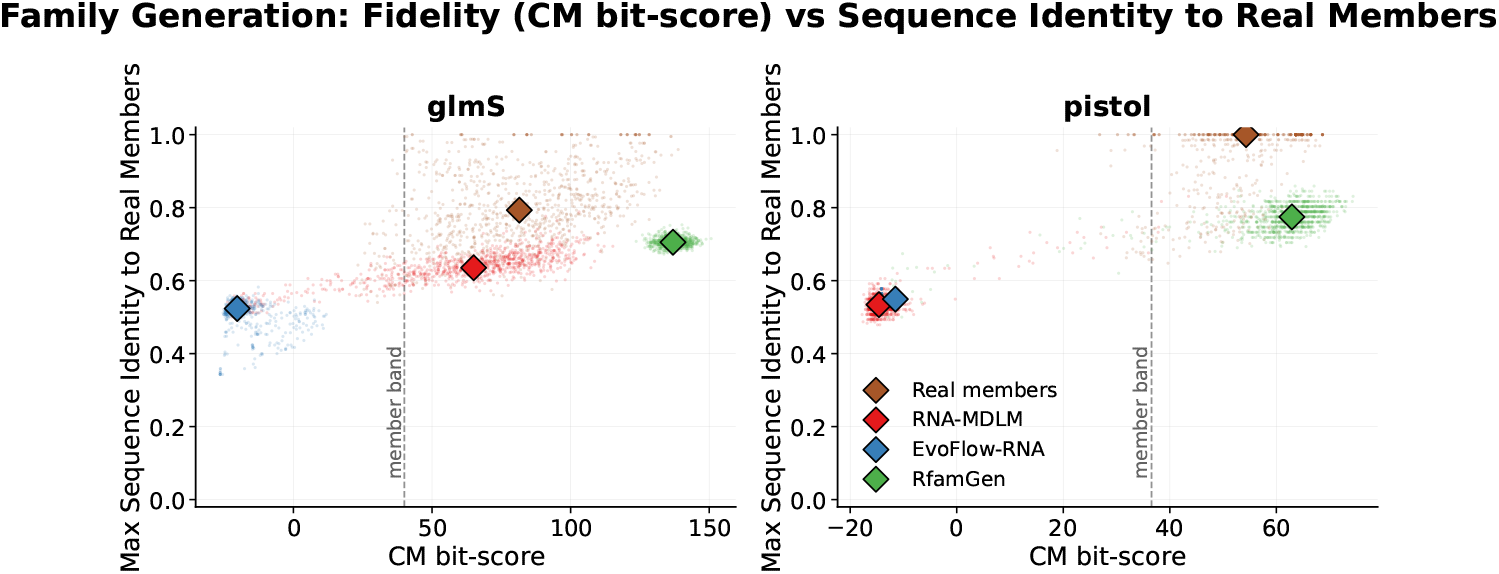
Within-family fidelity (CM bit-score, *x*) versus identity to real members (*y*; lower identity means more novel) for glmS (left) and Pistol (right). Genuine generation lies to the right of the member band (in-family) while staying low on identity (novel). RNA-MDLM (class-conditioned) reaches the glmS band at the lowest identity of any in-family model; RfamGen (per-family) is high-fidelity but sits at higher identity; EvoFlow-RNA (unconditional) reaches neither family.

**Figure 10:**
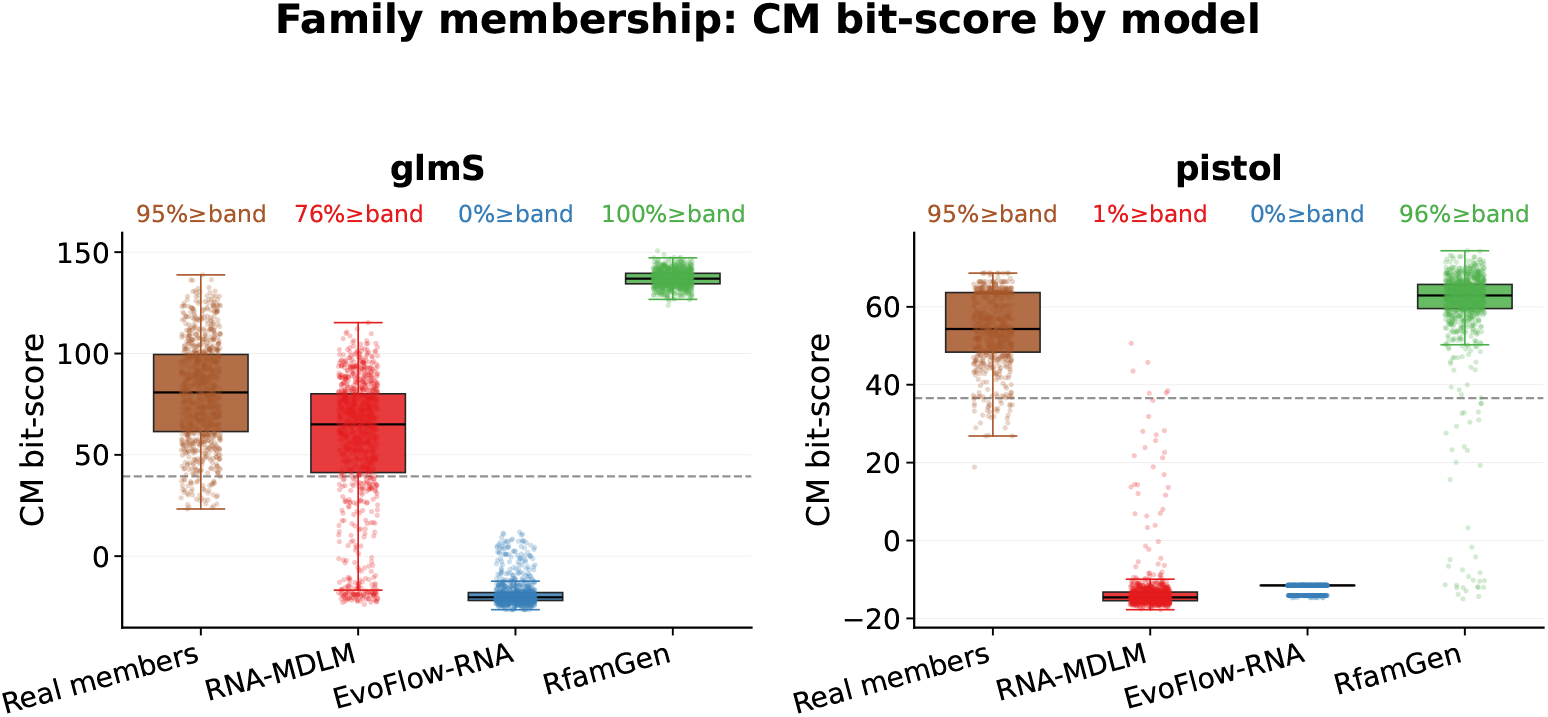
Per-model CM bit-score distributions for glmS and Pistol, with the fraction of each model’s sequences reaching the real-member band annotated. RNA-MDLM is class-conditioned; RfamGen is per-family; EvoFlow-RNA is unconditional.

**Figure 11:**
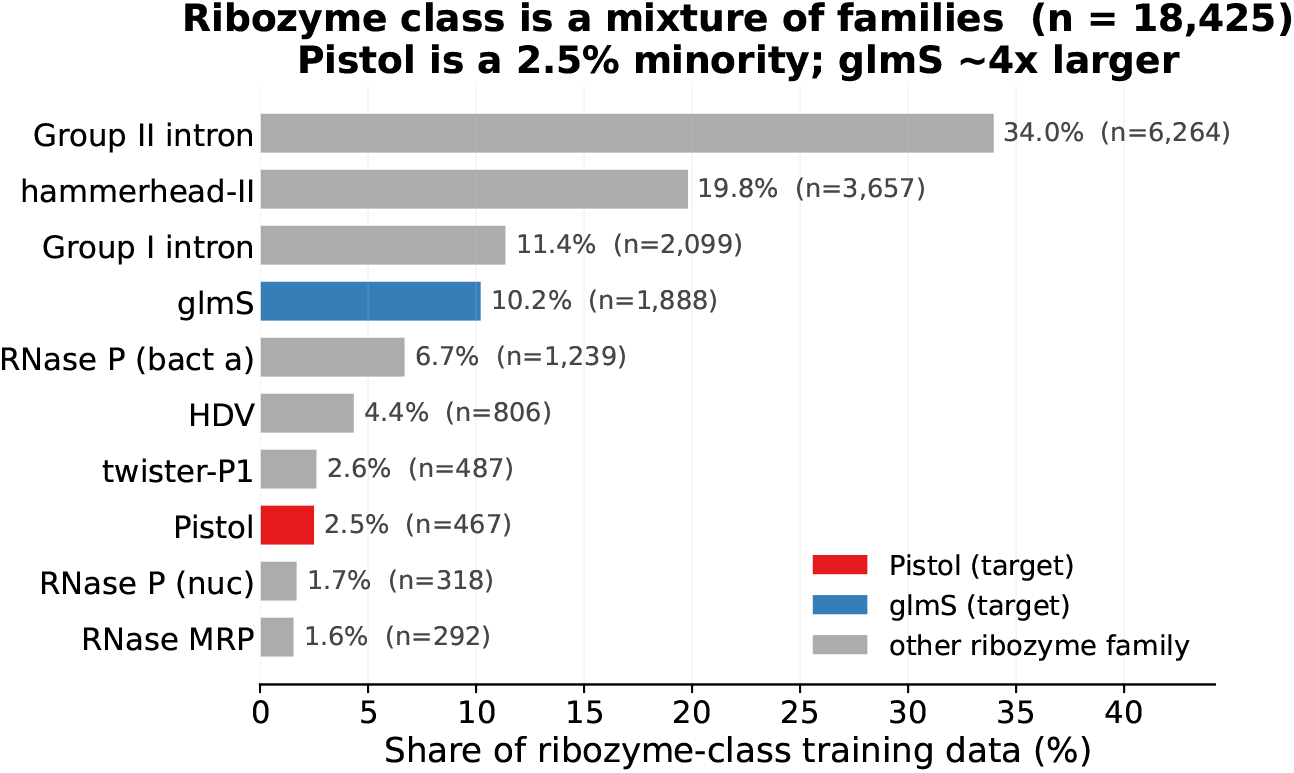
Family composition of the 18,425 ribozyme-class training sequences. Pistol (2.5%) is roughly 4*×* rarer than glmS (10.2%).

**Figure 12:**
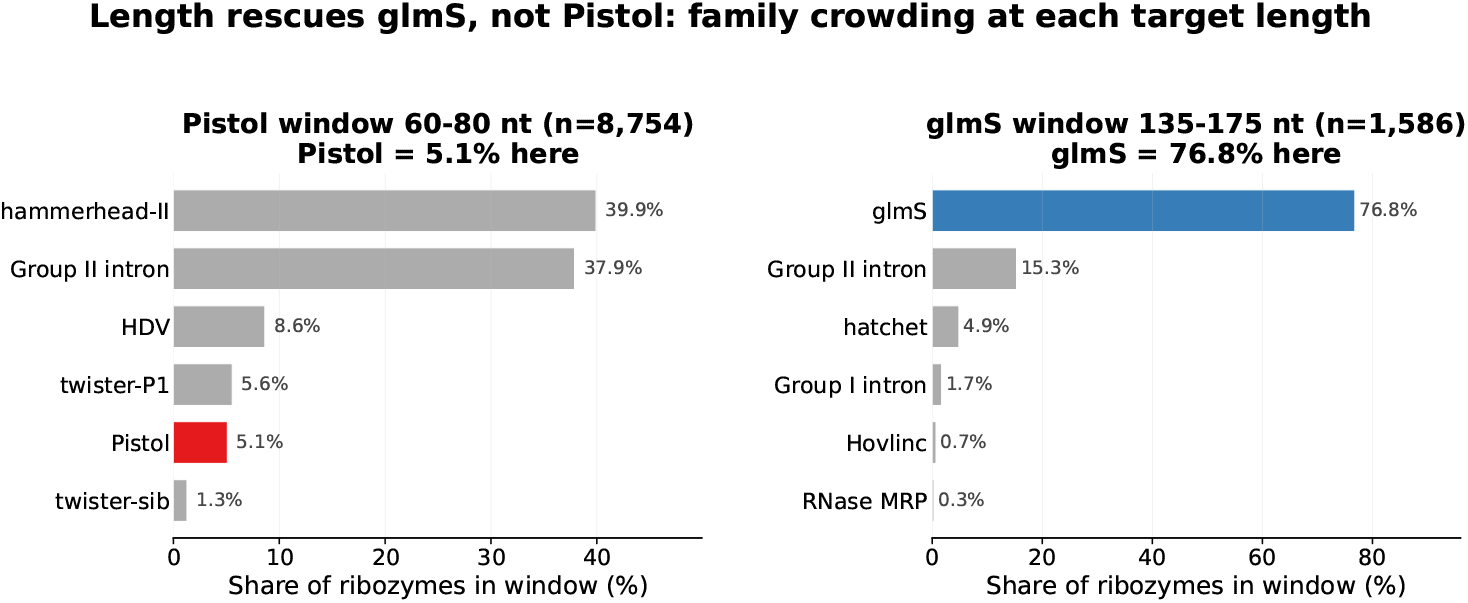
Family share of the ribozyme class at each target length. At Pistol’s 60–80 nt window Pistol is only 5.1% (hammerhead-II and Group II introns dominate), whereas at glmS’s 135–175 nt window glmS is 76.8%. Length accidentally isolates glmS but not Pistol.

**Figure 13:**
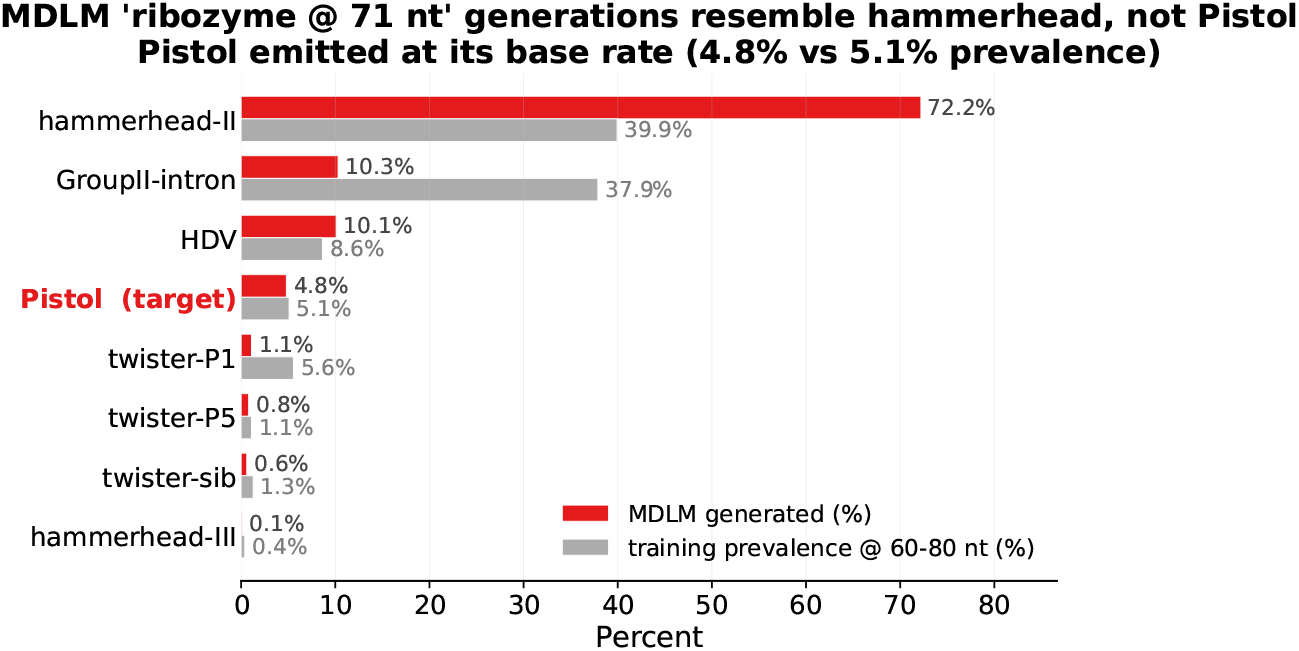
Family attribution of 1000 RNA-MDLM *ribozyme @ 71 nt* generations: 72.2% resemble hammerhead-II and 4.8% Pistol (near its 5.1% training prevalence). A positive control with real Pistol sequences as input attributes 100% to Pistol.

### A.11 RePaint Games: Per-game Recovery and Figures

**Figure 14:**
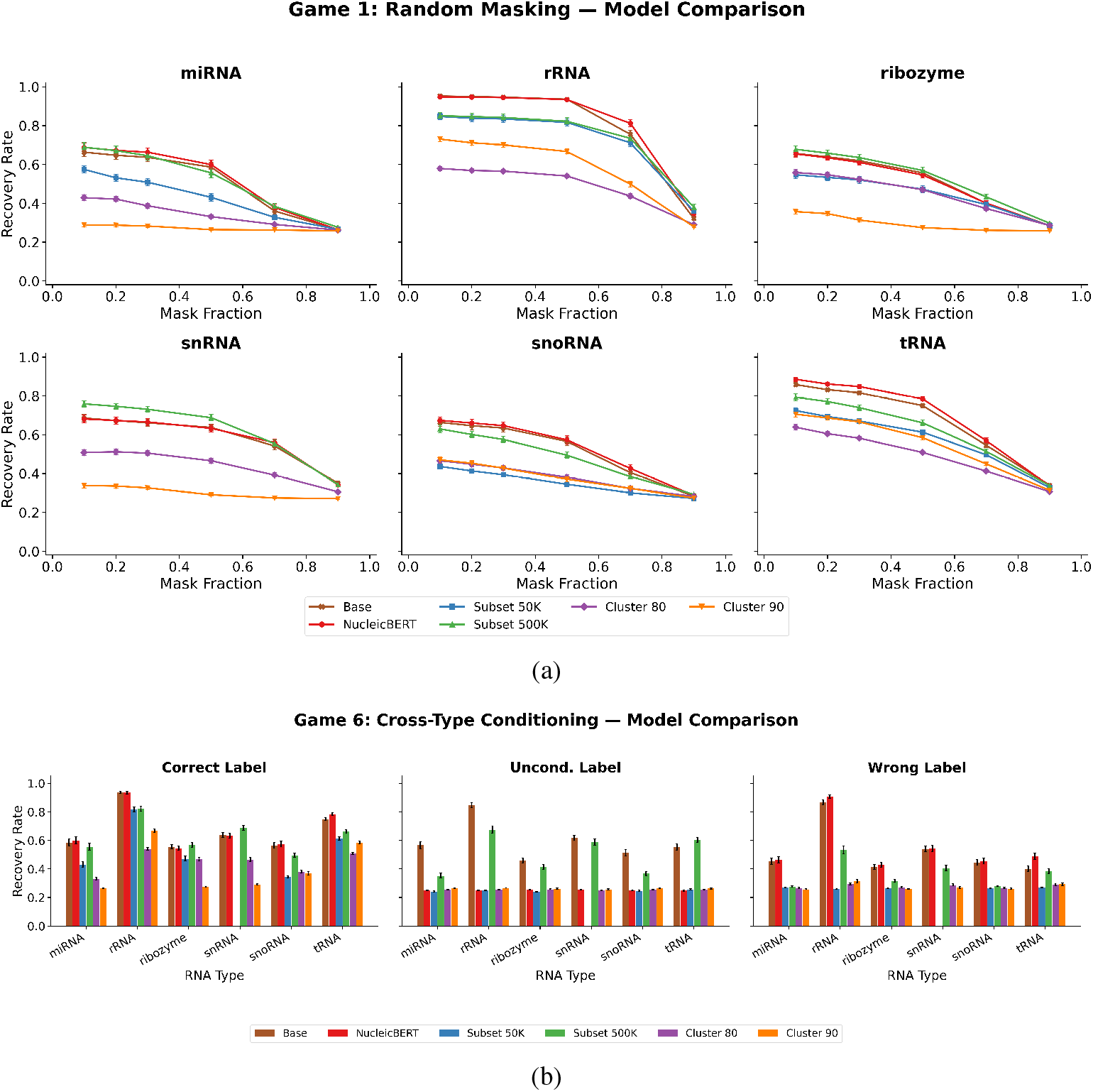
REPAINT GAMES recovery by RNA type and variant. (a) Game 1, random masking (recovery vs. masking fraction). (b) Game 7, structure preservation (closing-strand recovery). Error bars are the standard error over sequences.

**Figure 15:**
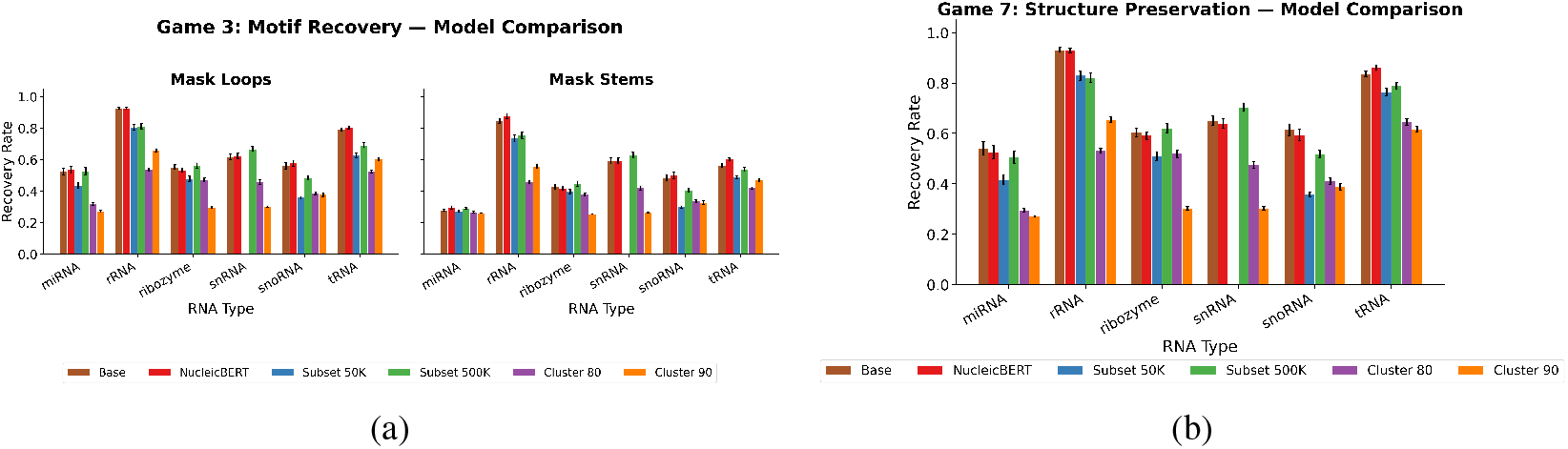
(a) Game 3, motif recovery (loops vs. stems). (b) Game 6, cross-type conditioning (correct, null, and wrong labels).

**Figure 16:**
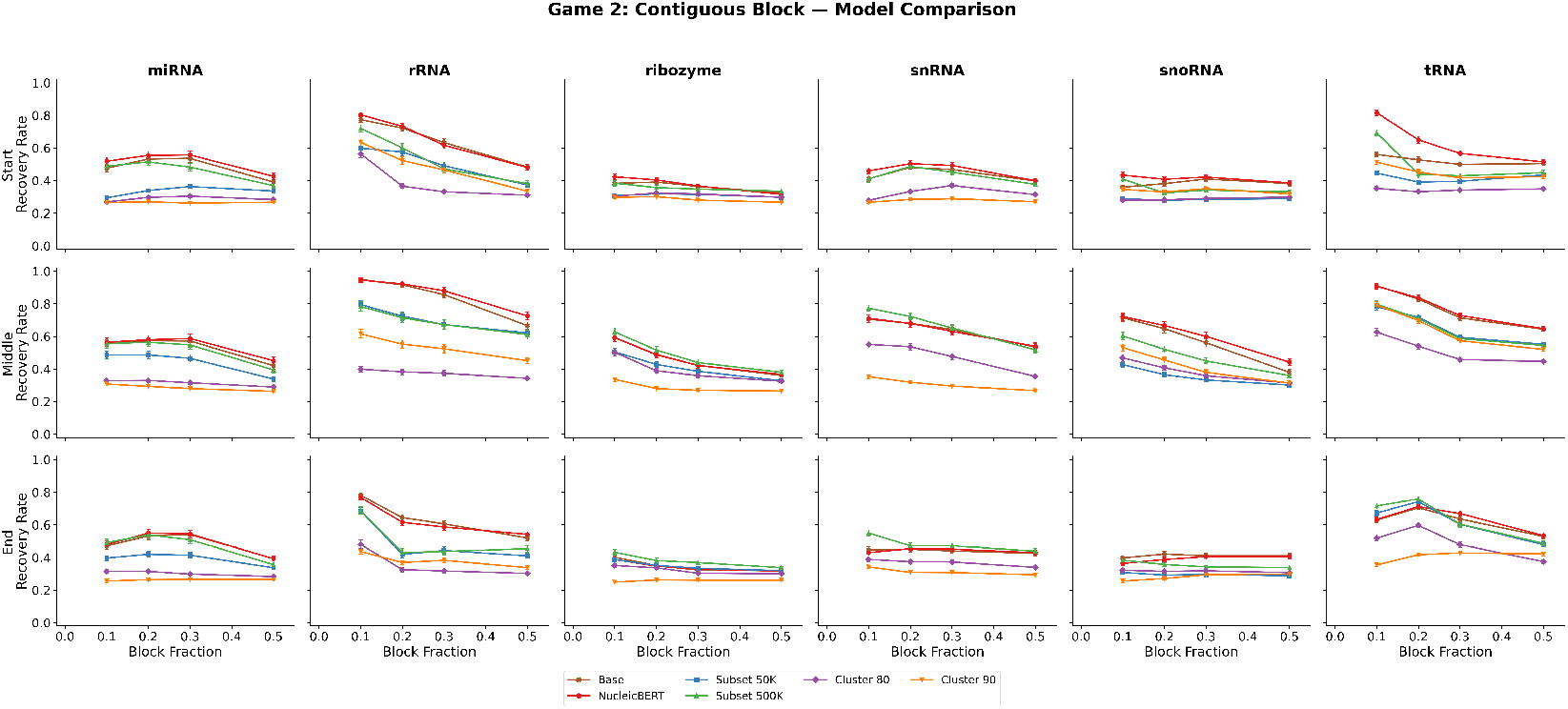
Game 2, contiguous block. Recovery vs. block fraction for masking at the 5’ start, middle, and 3’ end, by RNA type and variant.

**Figure 17:**
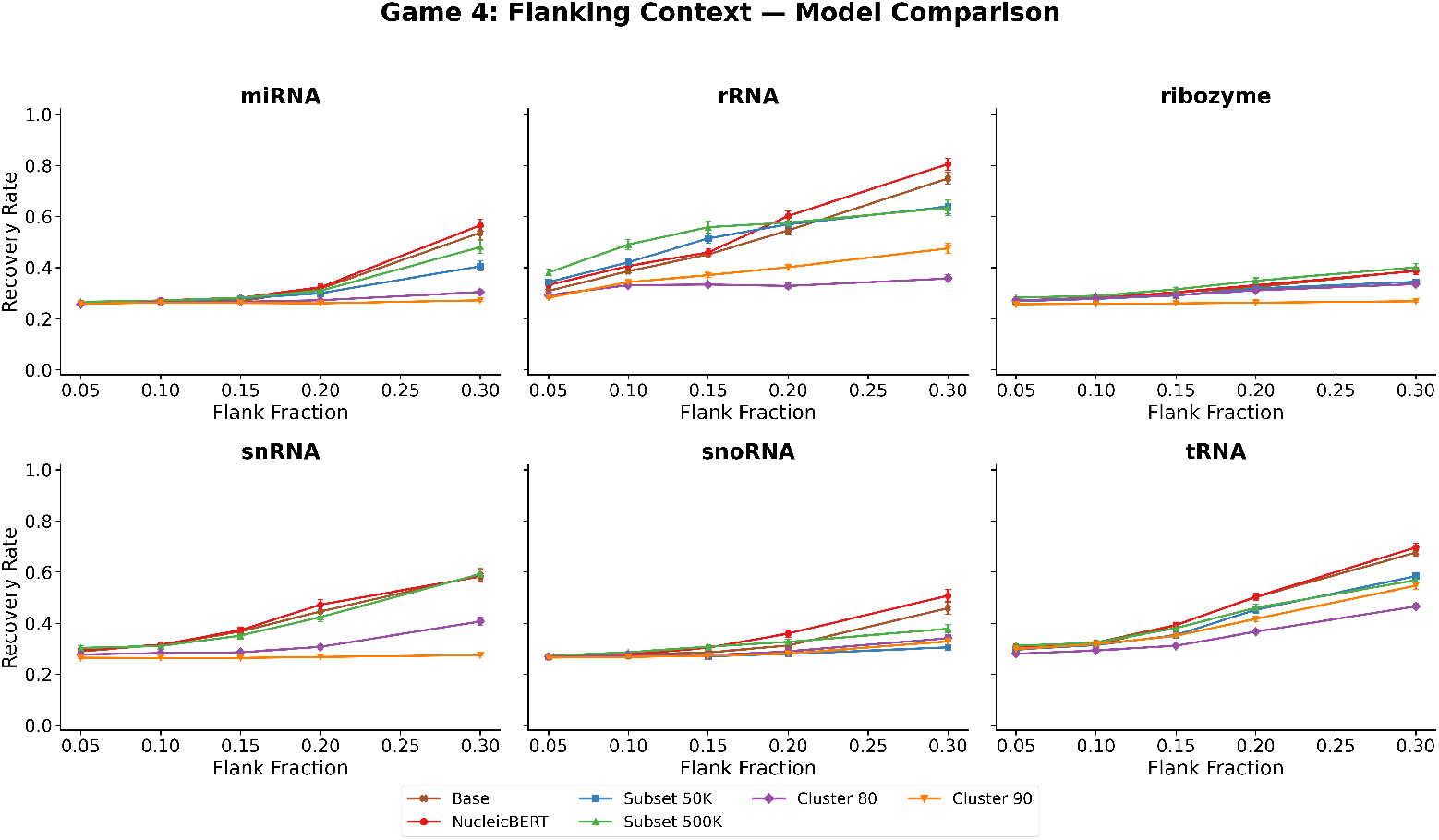
Game 4, flanking context. Recovery of the masked interior vs. the fraction of terminal flanking context retained.

**Figure 18:**
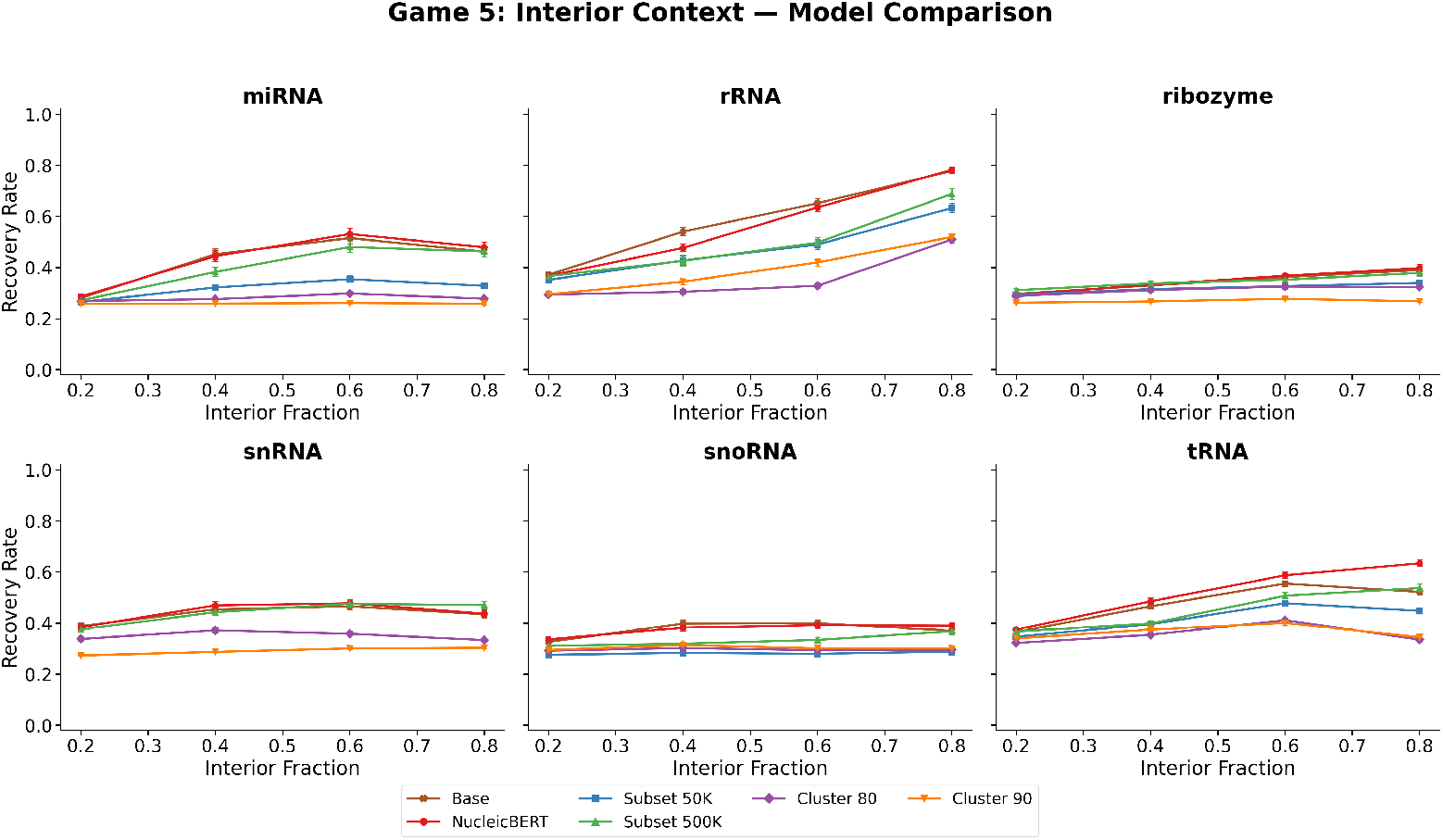
Game 5, interior context. Recovery of the masked ends vs. the fraction of interior context retained.

**Table 7:**
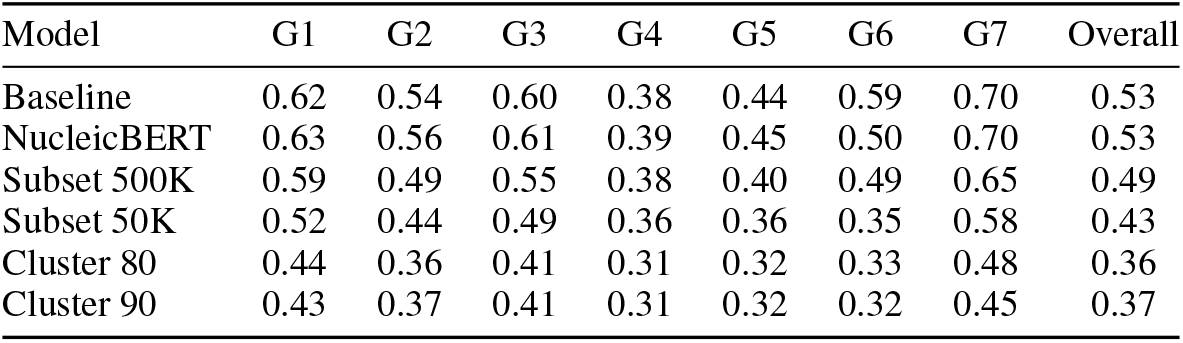
REPAINT GAMES recovery by variant and game, pooled over the five classes present in every subset. Overall pools all masked positions and is not the row mean (G6 = cross-type conditioning, G7 = structure preservation). The G6 column pools recovery over the correct, null, and wrong-label conditions, so a higher value there can reflect a variant *ignoring* the label rather than steering on it; the correct-vs-wrong steering gap is reported in Table 8.

| Model | G1 | G2 | G3 | G4 | G5 | G6 | G7 | Overall |
| --- | --- | --- | --- | --- | --- | --- | --- | --- |
| Baseline | 0.62 | 0.54 | 0.60 | 0.38 | 0.44 | 0.59 | 0.70 | 0.53 |
| NucleicBERT | 0.63 | 0.56 | 0.61 | 0.39 | 0.45 | 0.50 | 0.70 | 0.53 |
| Subset 500K | 0.59 | 0.49 | 0.55 | 0.38 | 0.40 | 0.49 | 0.65 | 0.49 |
| Subset 50K | 0.52 | 0.44 | 0.49 | 0.36 | 0.36 | 0.35 | 0.58 | 0.43 |
| Cluster 80 | 0.44 | 0.36 | 0.41 | 0.31 | 0.32 | 0.33 | 0.48 | 0.36 |
| Cluster 90 | 0.43 | 0.37 | 0.41 | 0.31 | 0.32 | 0.32 | 0.45 | 0.37 |

**Table 8:**
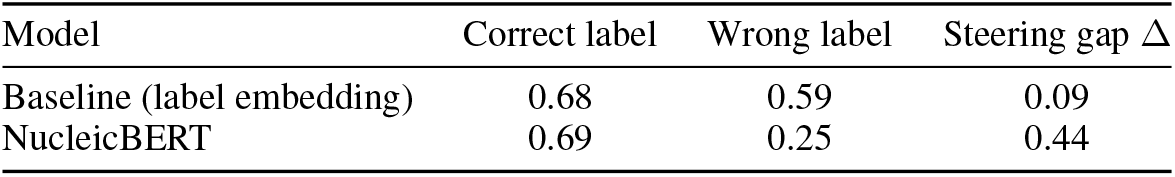
Cross-type steering (G6): recovery under the correct label versus the wrong label, and the gap Δ between them, for the two full-data conditioning schemes. A large Δ means the class label genuinely drives generation; a small Δ means the model recovers from sequence context and largely ignores the label. NucleicBERT conditioning steers far more strongly than the label embedding.

| Model | Correct label | Wrong label | Steering gap $\Delta$ |
| --- | --- | --- | --- |
| Baseline (label embedding) | 0.68 | 0.59 | 0.09 |
| NucleicBERT | 0.69 | 0.25 | 0.44 |

### A.12 EvoFlow-RNA Inpainting Comparison

This section reports the per-type REPAINT GAMES curves behind Table 2, one figure per game, for RNA-MDLM (Baseline and NucleicBERT) versus EvoFlow-RNA under the same masked sequences.

**Figure 19:**
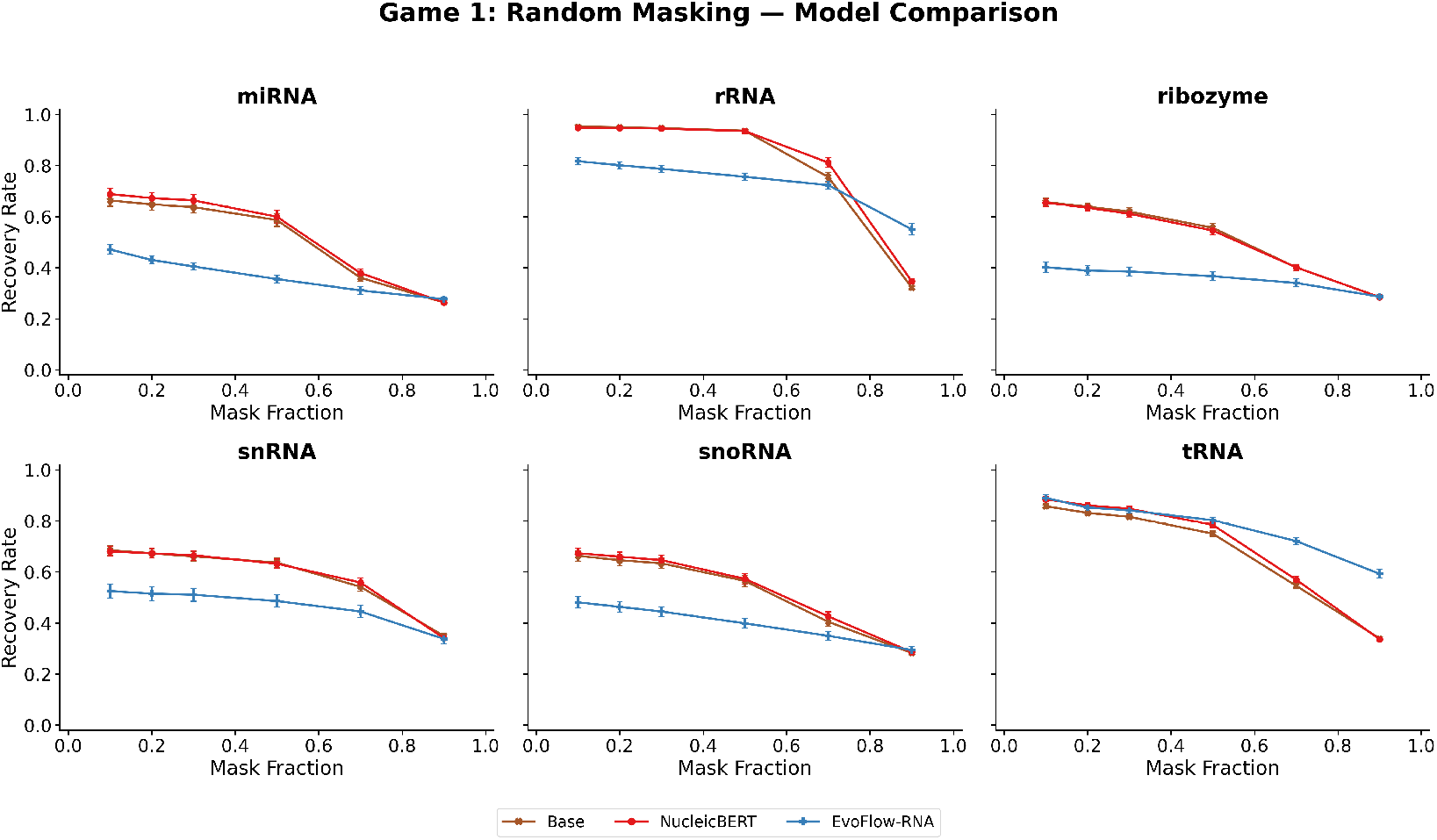
Game 1, random masking: per-type recovery vs. masking fraction for RNA-MDLM (Baseline and NucleicBERT) versus EvoFlow-RNA.

**Figure 20:**
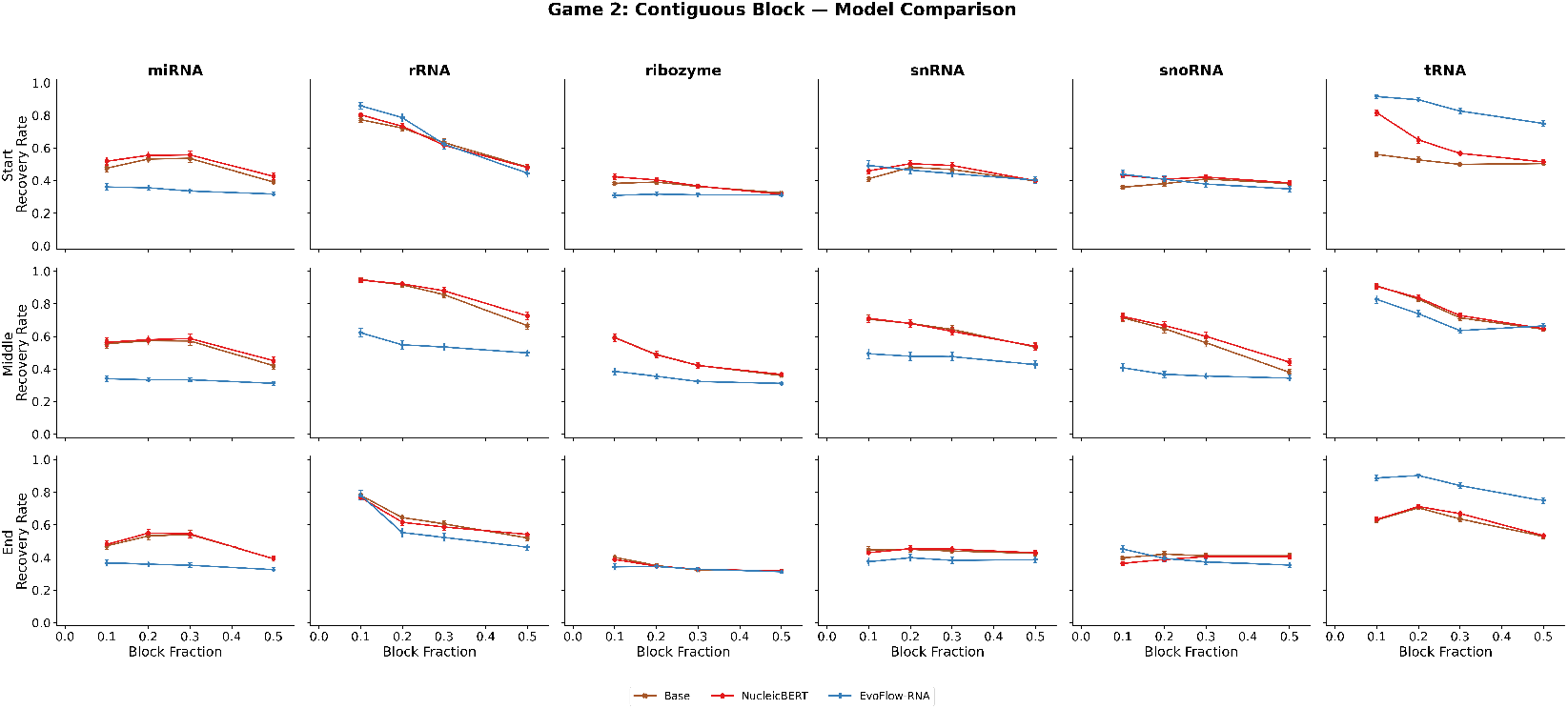
Game 2, contiguous block: per-type recovery vs. block fraction for masking at the 5’ start, middle, and 3’ end (rows), for RNA-MDLM versus EvoFlow-RNA.

**Figure 21:**
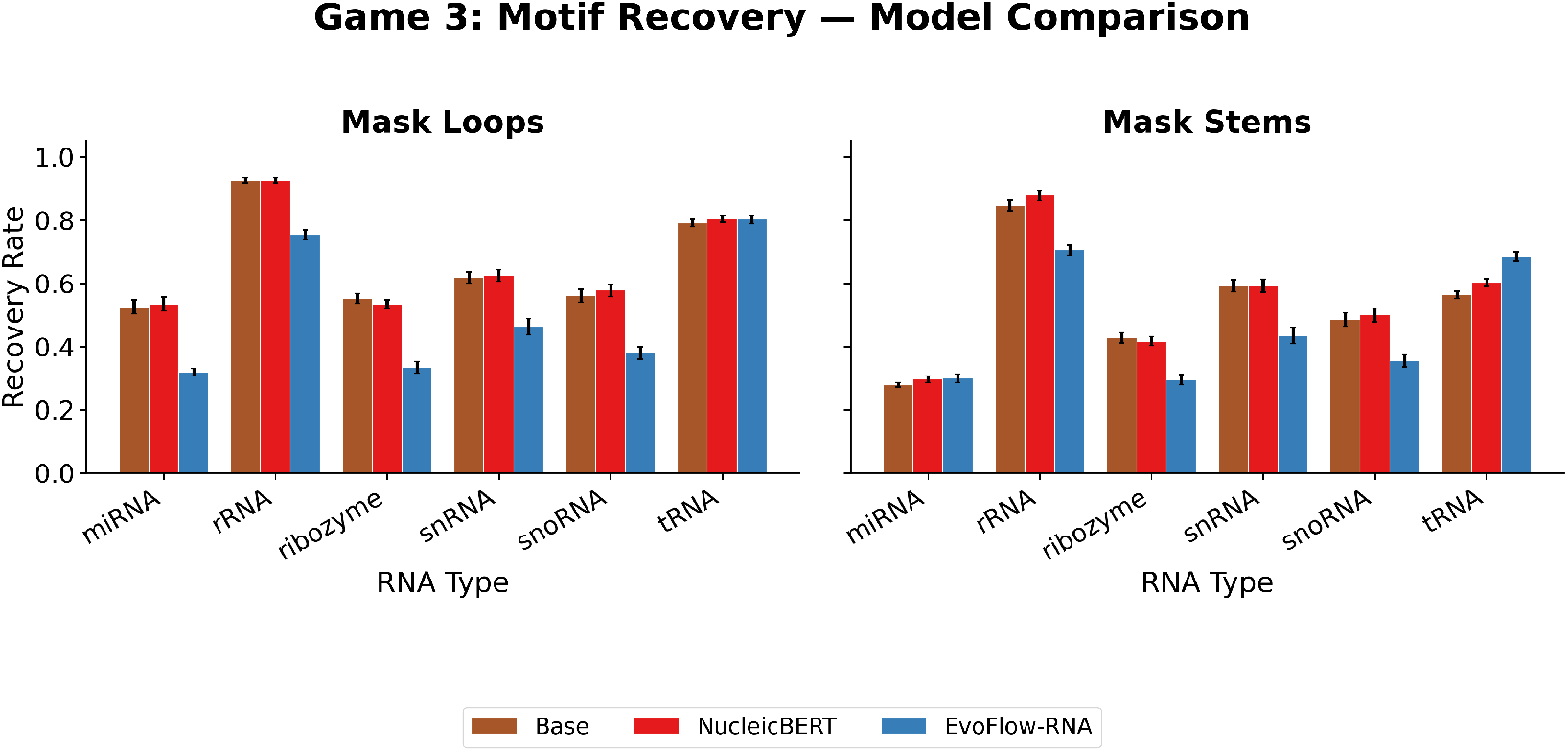
Game 3, motif recovery: per-type recovery when masking all loop positions or all stem positions, for RNA-MDLM versus EvoFlow-RNA.

**Figure 22:**
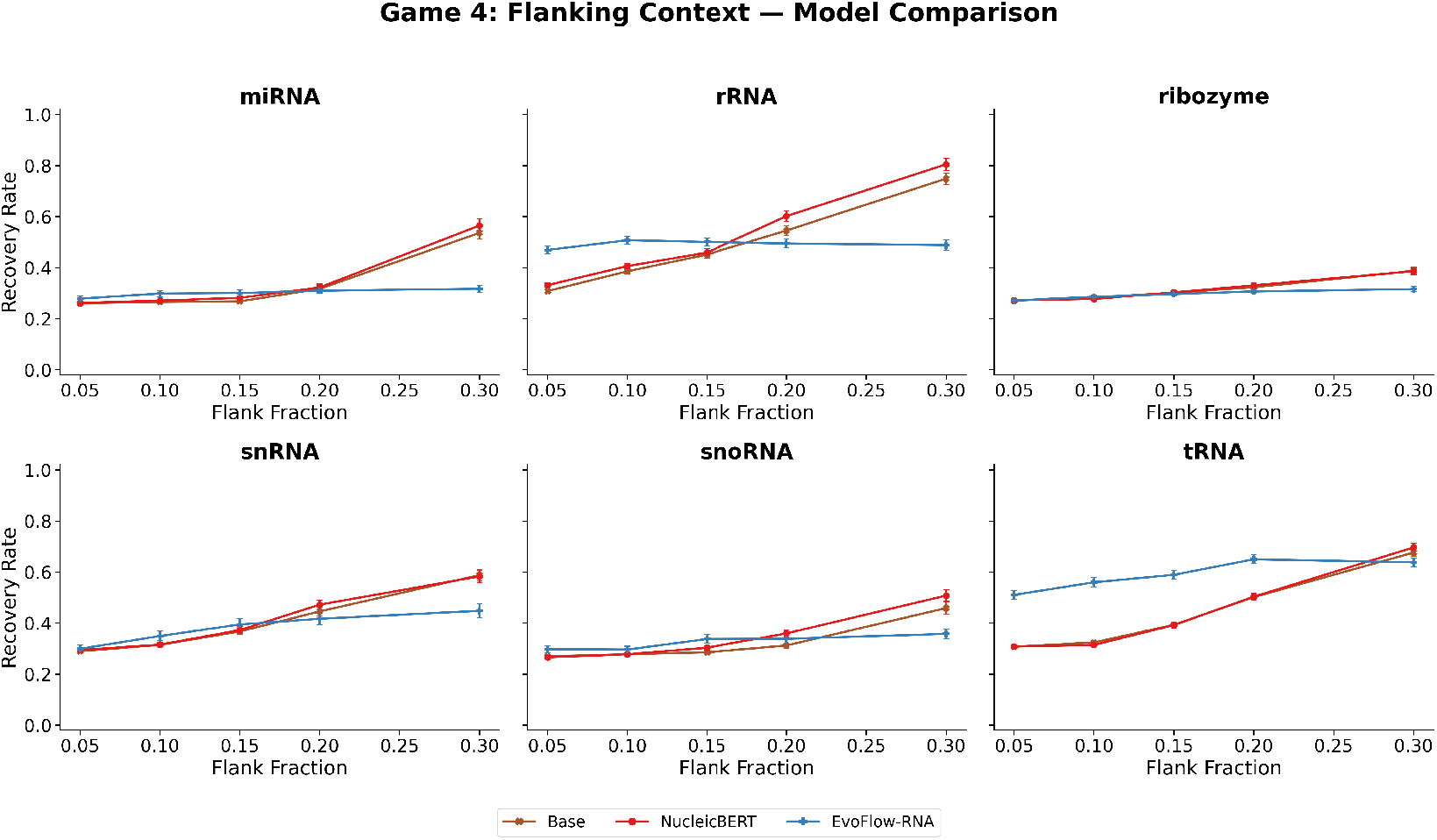
Game 4, flanking context: per-type recovery of the masked interior vs. the fraction of terminal flanking context retained, for RNA-MDLM versus EvoFlow-RNA.

**Figure 23:**
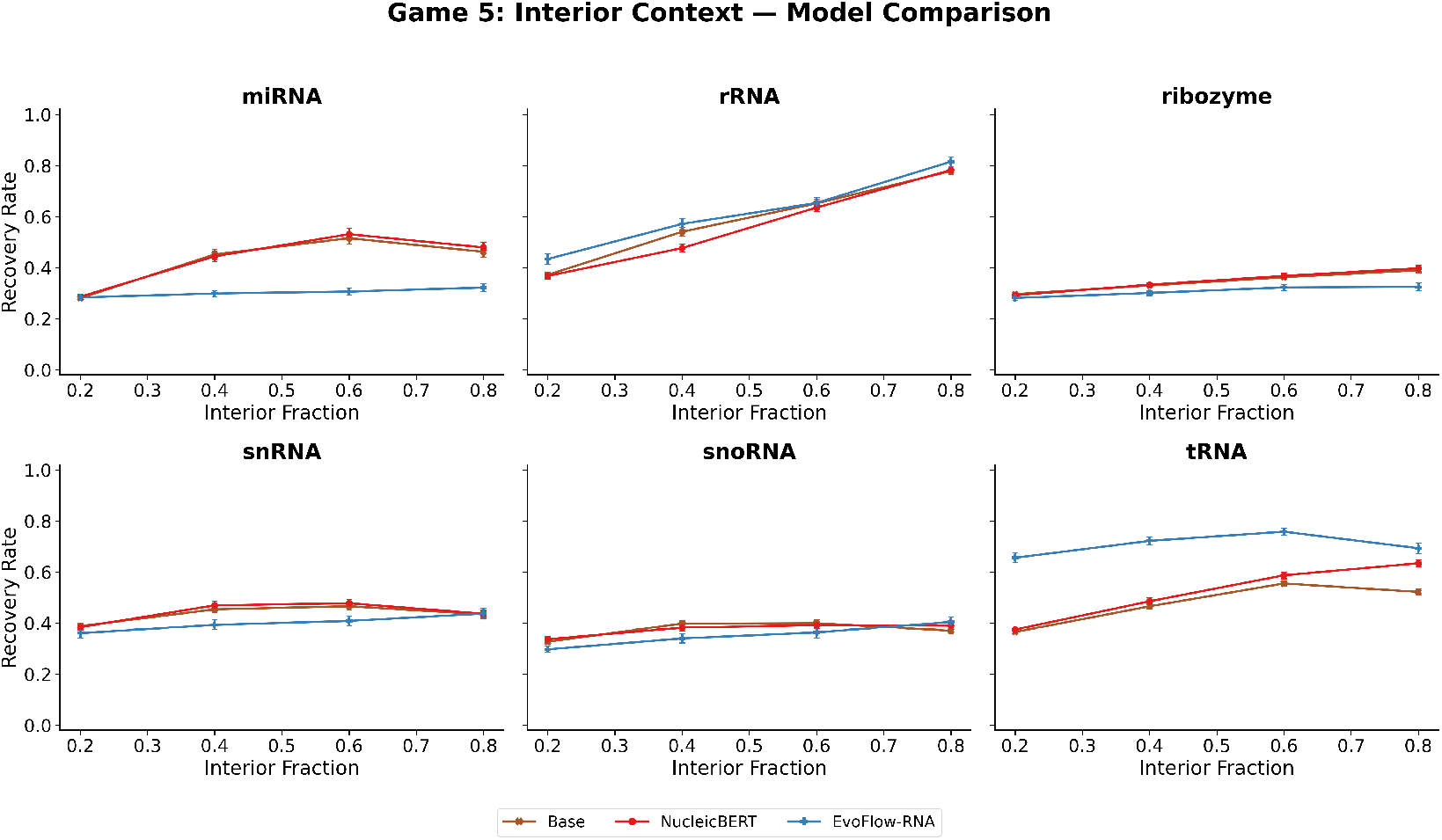
Game 5, interior context: per-type recovery of the masked ends vs. the fraction of interior context retained, for RNA-MDLM versus EvoFlow-RNA.

**Figure 24:**
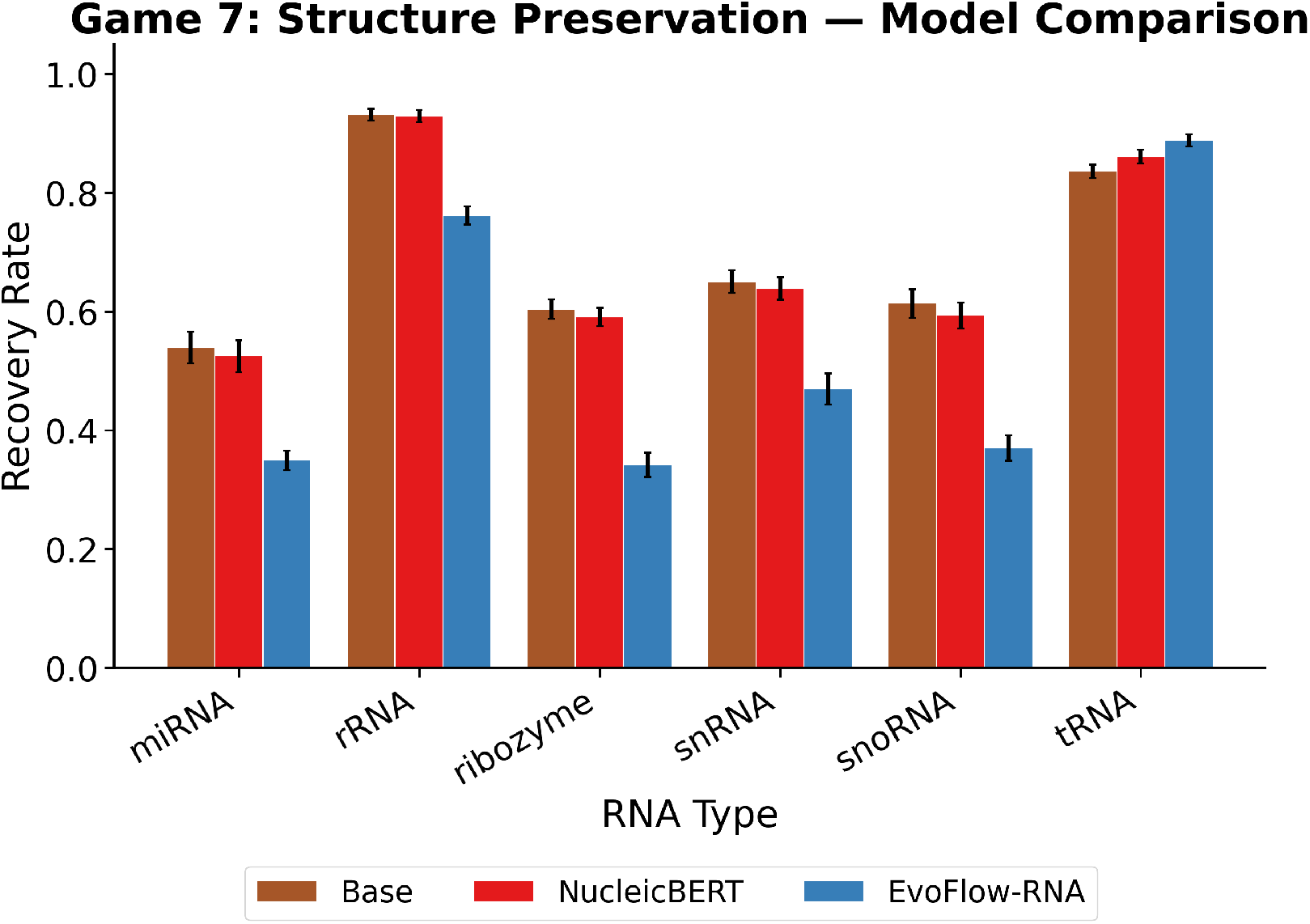
Game 7, structure preservation: per-type closing-strand recovery for RNA-MDLM versus EvoFlow-RNA.

## Notes

### Competing Interest Statement

The authors have declared no competing interest.

### Funding

- Helmholtz Association of German Research Centres

## References

1. Jacob Austin, Daniel D. Johnson, Jonathan Ho, Daniel Tarlow, and Rianne van den Berg. Structured Denoising Diffusion Models in Discrete State-Spaces. In Advances in Neural Information Processing Systems, volume 34, pp. 17981–17993. Curran Associates, Inc., 2021.

2. Pavel Avdeyev, Chenlai Shi, Yuhao Tan, Kseniia Dudnyk, and Jian Zhou. Dirichlet Diffusion Score Model for Biological Sequence Generation. In Proceedings of the 40th International Conference on Machine Learning, pp. 1276–1301. PMLR, July 2023.

3. David P. Bartel. Metazoan MicroRNAs. Cell, 173(1):20–51, March 2018. ISSN 0092-8674. doi: 10.1016/j.cell.2018.03.006.

4. Thomas R. Cech and Joan A. Steitz. The Noncoding RNA Revolution—Trashing Old Rules to Forge New Ones. Cell, 157(1):77–94, March 2014. ISSN 0092-8674. doi: 10.1016/j.cell.2014.03.008.

5. Tulsi Ram Damase, Roman Sukhovershin, Christian Boada, Francesca Taraballi, Roderic I. Pettigrew, and John P. Cooke. The Limitless Future of RNA Therapeutics. Frontiers in Bioengineering and Biotechnology, 9, March 2021. ISSN 2296-4185. doi: 10.3389/fbioe.2021.628137.

6. Tri Dao, Daniel Y. Fu, Stefano Ermon, Atri Rudra, and Christopher Ré. FLASHATTENTION: Fast and memory-efficient exact attention with IO-awareness. In Proceedings of the 36th International Conference on Neural Information Processing Systems, NIPS ’22, pp. 16344–16359, Red Hook, NY, USA, November 2022. Curran Associates Inc. ISBN 978-1-7138-7108-8.

7. J. Dauparas, I. Anishchenko, N. Bennett, H. Bai, R. J. Ragotte, L. F. Milles, B. I. M. Wicky, A. Courbet, R. J. de Haas, N. Bethel, P. J. Y. Leung, T. F. Huddy, S. Pellock, D. Tischer, F. Chan, B. Koepnick, H. Nguyen, A. Kang, B. Sankaran, A. K. Bera, N. P. King, and D. Baker. Robust deep learning–based protein sequence design using ProteinMPNN. Science, 378(6615):49–56, October 2022. doi: 10.1126/science.add2187.

8. Limin Fu, Beifang Niu, Zhengwei Zhu, Sitao Wu, and Weizhong Li. CD-HIT: Accelerated for clustering the next-generation sequencing data. Bioinformatics, 28(23):3150–3152, December 2012. ISSN 1367-4803. doi: 10.1093/bioinformatics/bts565.

9. Jonathan Ho and Tim Salimans. Classifier-Free Diffusion Guidance. In NeurIPS 2021 Workshop on Deep Generative Models and Downstream Applications, December 2021.

10. I. L. Hofacker, W. Fontana, P. F. Stadler, L. S. Bonhoeffer, M. Tacker, and P. Schuster. Fast folding and comparison of RNA secondary structures. Monatshefte für Chemie / Chemical Monthly, 125 (2):167–188, February 1994. ISSN 1434-4475. doi: 10.1007/BF00818163.

11. Han Huang, Ziqian Lin, Dongchen He, Liang Hong, and Yu Li. RiboDiffusion: Tertiary structurebased RNA inverse folding with generative diffusion models. Bioinformatics, 40(Supplement 1): i347–i356, July 2024a. ISSN 1367-4811. doi: 10.1093/bioinformatics/btae259.

12. Kaixuan Huang, Yukang Yang, Kaidi Fu, Yanyi Chu, Le Cong, and Mengdi Wang. Latent Diffusion Models for Controllable RNA Sequence Generation, October 2024b.

13. Yanjie Huang, Guangye Lv, Anyue Cheng, Wei Xie, Mengyan Chen, Xinyi Ma, Yijun Huang, Yueyang Tang, Qingya Shi, Zining Wang, Junxi Wang, Yunpeng Xia, Lu Zhao, Yifang Cai, Jack X. Chen, and Shuangjia Zheng. A Long-Context Generative Foundation Model Deciphers RNA Design Principles, March 2026. ISSN 2692-8205.

14. John B. Ingraham, Max Baranov, Zak Costello, Karl W. Barber, Wujie Wang, Ahmed Ismail, Vincent Frappier, Dana M. Lord, Christopher Ng-Thow-Hing, Erik R. Van Vlack, Shan Tie, Vincent Xue, Sarah C. Cowles, Alan Leung, João V. Rodrigues, Claudio L. Morales-Perez, Alex M. Ayoub, Robin Green, Katherine Puentes, Frank Oplinger, Nishant V. Panwar, Fritz Obermeyer, Adam R. Root, Andrew L. Beam, Frank J. Poelwijk, and Gevorg Grigoryan. Illuminating protein space with a programmable generative model. Nature, 623(7989):1070–1078, November 2023. ISSN 1476-4687. doi: 10.1038/s41586-023-06728-8.

15. Chaitanya K. Joshi, Arian R. Jamasb, Ramon Viñas, Charles Harris, Simon V. Mathis, Alex Morehead, Rishabh Anand, and Pietro Liò. gRNAde: Geometric Deep Learning for 3D RNA inverse design, February 2025.

16. Ioanna Kalvari, Eric P Nawrocki, Nancy Ontiveros-Palacios, Joanna Argasinska, Kevin Lamkiewicz, Manja Marz, Sam Griffiths-Jones, Claire Toffano-Nioche, Daniel Gautheret, Zasha Weinberg, Elena Rivas, Sean R Eddy, Robert D Finn, Alex Bateman, and Anton I Petrov. Rfam 14: Expanded coverage of metagenomic, viral and microRNA families. Nucleic Acids Research, 49(D1):D192–D200, January 2021. ISSN 0305-1048. doi: 10.1093/nar/gkaa1047.

17. Sizhen Li, Saeed Moayedpour, Ruijiang Li, Michael Bailey, Saleh Riahi, Lorenzo Kogler-Anele, Milad Miladi, Jacob Miner, Fabien Pertuy, Dinghai Zheng, Jun Wang, Akshay Balsubramani, Khang Tran, Minnie Zacharia, Monica Wu, Xiaobo Gu, Ryan Clinton, Carla Asquith, Joseph Skaleski, Lianne Boeglin, Sudha Chivukula, Anusha Dias, Tod Strugnell, Fernando Ulloa Montoya, Vikram Agarwal, Ziv Bar-Joseph, and Sven Jager. CodonBERT large language model for mRNA vaccines. Genome Research, 34(7):1027–1035, August 2024. ISSN 1088-9051, 1549-5469. doi: 10.1101/gr.278870.123.

18. Ronny Lorenz, Stephan H. Bernhart, Christian Höner zu Siederdissen, Hakim Tafer, Christoph Flamm, Peter F. Stadler, and Ivo L. Hofacker. ViennaRNA Package 2.0. Algorithms for Molecular Biology, 6(1):26, November 2011. ISSN 1748-7188. doi: 10.1186/1748-7188-6-26.

19. Aaron Lou, Chenlin Meng, and Stefano Ermon. Discrete diffusion modeling by estimating the ratios of the data distribution. In Proceedings of the 41st International Conference on Machine Learning, volume 235 of *ICML’24*, pp. 32819–32848, Vienna, Austria, July 2024. JMLR.org.

20. Andreas Lugmayr, Martin Danelljan, Andres Romero, Fisher Yu, Radu Timofte, and Luc Van Gool. RePaint: Inpainting using Denoising Diffusion Probabilistic Models. In 2022 IEEE/CVF Conference on Computer Vision and Pattern Recognition (CVPR), pp. 11451–11461, June 2022. doi: 10.1109/CVPR52688.2022.01117.

21. Eric P. Nawrocki and Sean R. Eddy. Infernal 1.1: 100-fold faster RNA homology searches. Bioinformatics, 29(22):2933–2935, November 2013. ISSN 1367-4803. doi: 10.1093/bioinformatics/btt509.

22. Sawan Patel, Fred Zhangzhi Peng, Keith Fraser, Pranam Chatterjee, and Sherwood Yao. EvoFlow-RNA: Generating and Representing non-coding RNA with a Language Model, February 2025.

23. William Peebles and Saining Xie. Scalable Diffusion Models with Transformers. In Proceedings of the IEEE/CVF International Conference on Computer Vision, pp. 4195–4205, 2023.

24. RNAcentral Consortium. RNAcentral 2021: Secondary structure integration, improved sequence search and new member databases. Nucleic Acids Research, 49(D1):D212–D220, January 2021. ISSN 0305-1048. doi: 10.1093/nar/gkaa921.

25. Ugur Sahin, Katalin Karikó, and Özlem Türeci. mRNA-based therapeutics — developing a new class of drugs. Nature Reviews Drug Discovery, 13(10):759–780, October 2014. ISSN 1474-1784. doi: 10.1038/nrd4278.

26. Subham Sekhar Sahoo, Marianne Arriola, Yair Schiff, Aaron Gokaslan, Edgar Marroquin, Justin T Chiu, Alexander Rush, and Volodymyr Kuleshov. Simple and Effective Masked Diffusion Language Models. In *Advances in Neural Information Processing Systems*, volume 37, pp. 130136– 130184. Curran Associates, Inc., 2024. doi: 10.52202/079017-4135.

27. Yair Schiff, Subham Sekhar Sahoo, Hao Phung, Guanghan Wang, Alexander Rush, Volodymyr Kuleshov, Hugo Dalla-Torre, Sam Boshar, Bernardo P. de Almeida, and Thomas Pierrot. Simple Guidance Mechanisms for Discrete Diffusion Models. International Conference on Learning Representations, 2025:44153–44198, April 2025.

28. Jianlin Su, Murtadha Ahmed, Yu Lu, Shengfeng Pan, Wen Bo, and Yunfeng Liu. RoFormer: Enhanced transformer with Rotary Position Embedding. Neurocomputing, 568:127063, February 2024. ISSN 0925-2312. doi: 10.1016/j.neucom.2023.127063.

29. Shunsuke Sumi, Michiaki Hamada, and Hirohide Saito. Deep generative design of RNA family sequences. Nature Methods, 21(3):435–443, March 2024. ISSN 1548-7105. doi: 10.1038/s41592-023-02148-8.

30. Utkarsh Upadhyay, Julian Herold, Markus Götz, and Alexander Schug. NucleicBERT interprets RNA sequence space through self-supervised language modelling. Nature Machine Intelligence, pp. 1–15, September 2026. ISSN 2522-5839. doi: 10.1038/s42256-026-01295-9.

31. Ashish Vaswani, Noam Shazeer, Niki Parmar, Jakob Uszkoreit, Llion Jones, Aidan N Gomez, Łukasz Kaiser, and Illia Polosukhin. Attention is All you Need. In Advances in Neural Information Processing Systems, volume 30. Curran Associates, Inc., 2017.

32. Joseph L. Watson, David Juergens, Nathaniel R. Bennett, Brian L. Trippe, Jason Yim, Helen E. Eisenach, Woody Ahern, Andrew J. Borst, Robert J. Ragotte, Lukas F. Milles, Basile I. M. Wicky, Nikita Hanikel, Samuel J. Pellock, Alexis Courbet, William Sheffler, Jue Wang, Preetham Venkatesh, Isaac Sappington, Susana Vázquez Torres, Anna Lauko, Valentin De Bortoli, Emile Mathieu, Sergey Ovchinnikov, Regina Barzilay, Tommi S. Jaakkola, Frank DiMaio, Minkyung Baek, and David Baker. De novo design of protein structure and function with RFdiffusion. Nature, 620(7976):1089–1100, August 2023. ISSN 1476-4687. doi: 10.1038/s41586-023-06415-8.

33. Yichong Zhao, Kenta Oono, Hiroki Takizawa, and Masaaki Kotera. GenerRNA: A generative pretrained language model for de novo RNA design. PLOS ONE, 19(10):e0310814, October 2024. ISSN 1932-6203. doi: 10.1371/journal.pone.0310814.

34. Lin Zheng, Jianbo Yuan, Lei Yu, and Lingpeng Kong. A Reparameterized Discrete Diffusion Model for Text Generation. In First Conference on Language Modeling, August 2024.

